# Information Flow in Planar Polarity

**DOI:** 10.1101/236836

**Authors:** Katherine H Fisher, David Strutt, Alexander G Fletcher

## Abstract

In developing tissues, sheets of cells become planar polarised, enabling coordination of cell behaviours. It has been suggested that ‘signalling’ of polarity information between cells may occur either bidirectionally or monodirectionally between the molecules Frizzled (Fz) and Van Gogh (Vang). Using computational modelling we find that both bidirectional and monodirectional signalling models reproduce known non-autonomous phenotypes derived from patches of mutant tissue of key molecules, but predict different phenotypes from double mutant tissue, which have previously given conflicting experimental results. Consequently, we re-examine experimental phenotypes in the *Drosophila* wing, concluding that signalling is most likely bidirectional. Our modelling suggests that bidirectional signalling can be mediated either *indirectly* via bidirectional feedbacks between asymmetric intercellular protein complexes, or *directly* via different affinities for protein binding in intercellular complexes, suggesting future avenues for investigation. Our findings offer insight into mechanisms of juxtacrine cell signalling and how tissue-scale properties emerge from individual cell behaviours.

## Introduction

### Planar polarity and patterning of the insect cuticle

In multicellular organisms, cells in a tissue frequently adopt a common polarity such that they are oriented in the same direction in the plane of the tissue. This is termed ‘planar polarity’ (or planar cell polarity [PCP]) and is necessary for morphogenesis, for instance ensuring cells within a group all move or intercalate along the same tissue axis. Furthermore, planar polarity is essential for tissue function, for example when motile cilia on the surface of an epithelium all adopt the same orientation and beat in the same direction (reviewed in Goodrich and Strutt, 2011; Devenport, 2014; Butler and Wallingford, 2017; Davey and Moens, 2017).

Cells within a tissue could each independently establish their planar polarity by reference to an external cue such as a gradient of an extracellular signalling molecule, biasing protein localisations to one or other side of a cell. Small biases could then be amplified through positive feedback to generate strong polarity (Tree et al., 2002; Amonlirdviman et al., 2005; Le Garrec et al., 2006; Abley et al., 2013; Warrington et al., 2017). However, variation in signal levels across the axis of a cell might be small and difficult to discriminate, leading to individual cells mispolarising. A solution is for cells to interact: comparing and coordinating polarity with their neighbours, thus establishing uniform polarity across a field of cells even in cases where the external graded signal is weak or noisy (Ma et al., 2003; Burak and Shraiman, 2009).

Evidence for cell-cell interactions during planar polarisation was provided by early transplantation experiments (Piepho, 1955; Locke, 1959) and later by direct manipulation of underlying genetic pathways in the fruit fly *Drosophila* (reviewed in Strutt, 2009). The best-studied pathway is known as the ‘core’ pathway (Goodrich and Strutt, 2011; Aw and Devenport, 2017), which shows clear evidence of cell-cell communication. However, some experimental results remain controversial, leading to uncertainty about the nature of such ‘signalling’.

### The core planar polarity pathway

The core pathway has six known protein components, which physically interact to form intercellular complexes at apicolateral cell junctions (Fig.1A). In planar polarised tissues, these complexes are asymmetrically distributed to opposite cell ends. In the developing *Drosophila* wing, the sevenpass transmembrane protein Fz localises to the distal side of cells (i.e. towards the tip of the wing) with the cytoplasmic proteins Dishevelled (Dsh) and Diego (Dgo), whereas the fourpass transmembrane protein Vang (also known as Strabismus [Stbm]) localises proximally (closest to the hinge or body of the fly) with the cytoplasmic protein Prickle (Pk). The sevenpass transmembrane atypical cadherin Flamingo (Fmi, also known as Starry Night [Stan]) forms intercellular homodimers and localises both proximally and distally, bridging the two halves of the complex (Strutt and Strutt, 2009) (Fig.1A). The asymmetric distribution of these proteins specifies the distal position from which an actin-rich hair, or trichome, emerges in each cell (Fig.1B).

**Figure 1.**
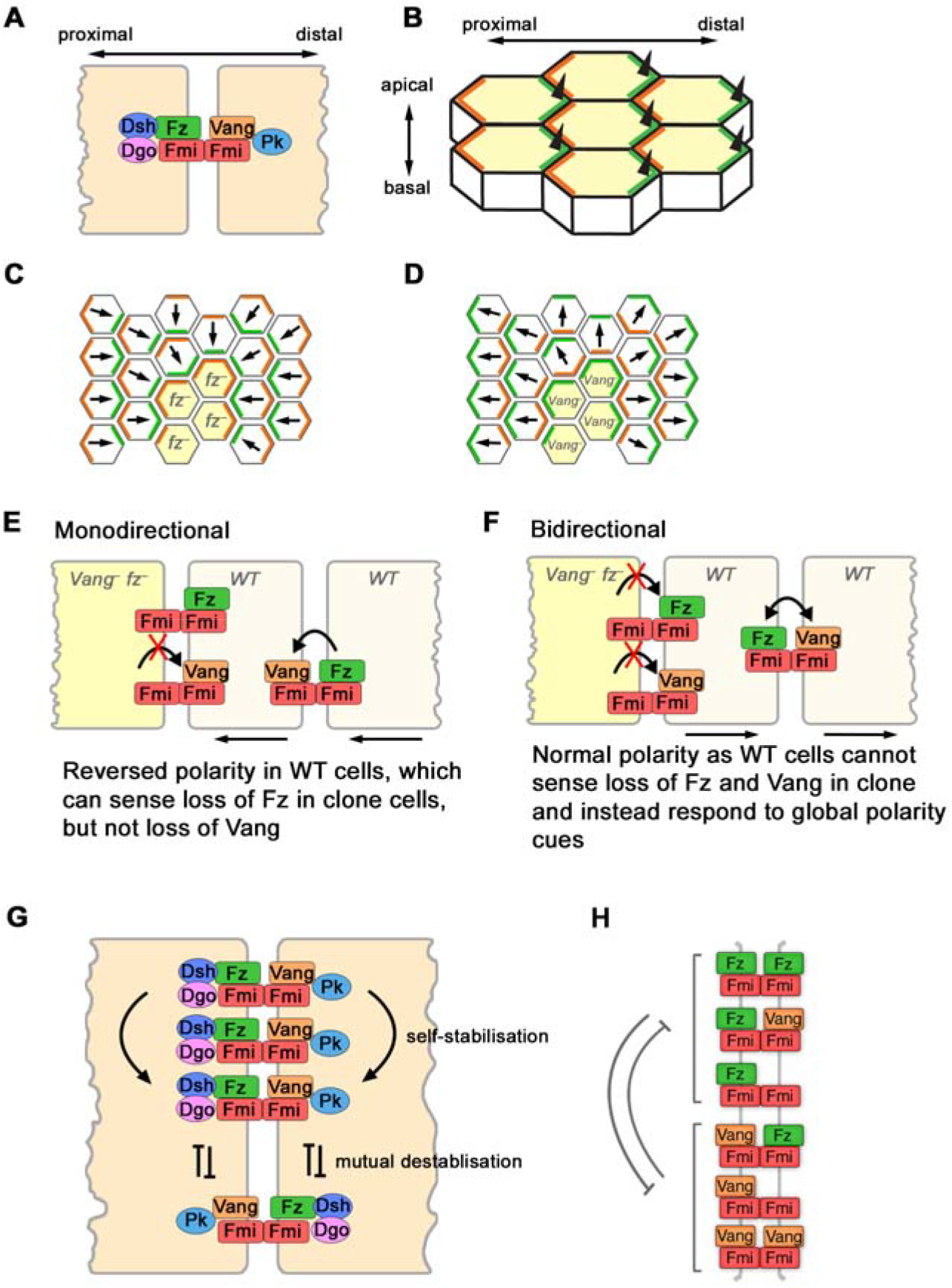
Directional signalling in planar polarity. (A) Diagram of the core planar polarity proteins localising to form an intercellular complex at a junction between two cells. (B) Diagram showing asymmetric localisation of Fz (green) and Vang (orange) in the developing *Drosophila* wing. Black triangles represent trichomes emerging at distal cell ends of the apical wing surface. (C,D) Diagrams of non-autonomous phenotypes of *fz*^-^ (C) and *Vang*^-^ (D) clones (yellow cells) in pupal wings. Arrows represent trichome orientation. At the edge of a *fz*^-^ clone, Vang within the clone cells localises to the boundary with wild-type neighbours, indicating it preferentially forms asymmetric complexes with Fz at cell-cell contacts. Similarly, Fz localises at the boundary of *Vang*^-^ clones. (E,F) Monodirectional (E) and bidirectional (F) signalling predictions. In monodirectional signalling (E), Vang acts as a ligand for Fz and can ‘sense’ its levels in neighbouring wild-type (WT) cells. *Vang*^-^ *fz*^-^ clones behave identically to *fz*^-^ clones, since no information on Vang levels is transmitted between cells. Cells distal to the clone have reversed polarity (black arrows). In bidirectional signalling (F), no information can be passed from *Vang*^-^ *fz*^-^ cells since both are required for sending and receiving the signal. Thus neighbouring wild-type cells are blind to the clone and polarise normally (black arrows). (G) Diagram of possible feedback interactions. Stabilising interactions occur locally between complexes of ‘like’ orientations, whereas destabilising interactions occur between complexes of ‘unlike’ orientations. (H) Diagram of destabilising feedbacks simulated between different possible Fz-and Vang-containing complexes localised at a junction between two cells in our computational modelling framework.

Strikingly, non-cell autonomous activity within this pathway is observed when patches of cells lacking Fz activity are juxtaposed to cells with Fz activity during wing development. The *fz* mutant cells modify the polarity of their neighbours, whose trichomes point towards the *fz* mutant tissue (Gubb and Garcia-Bellido, 1982; Vinson and Adler, 1987). This is accompanied by relocalisation of the core proteins parallel to the clone boundary (Fig.1C) (Usui et al., 1999; Strutt, 2001; Bastock et al., 2003). Similarly, clones of cells lacking the activity of *Vang* alter the polarity of neighbouring cells, in this case causing their trichomes to point away from the mutant tissue (Fig.1D) (Taylor et al., 1998).

The reorganisation of polarity around clones of cells lacking Fz and Vang, and their colocalisation on apposing junctions, suggests direct roles for these proteins in cell-cell communication. However, loss of Fmi in clones results in loss of all other core components from junctions (including Fz and Vang) (Strutt, 2001; Feiguin et al., 2001; Axelrod, 2001; Tree et al., 2002; Bastock et al., 2003), but does not cause significant repolarisation of neighbouring cells (Usui et al., 1999; Chae et al., 1999). Furthermore, cells with altered Fz or Vang activity, but lacking Fmi, can no longer repolarise their neighbours, and cells lacking Fmi cannot be repolarised by neighbours with altered Fz or Vang activity (Lawrence et al., 2004; Strutt and Strutt, 2007; Chen et al., 2008). These data support the view that Fz-Fmi complexes in each cell interact with Fmi-Vang complexes in neighbouring cells, and these polarised molecular bridges are the conduits for cell-cell transmission of polarity information.

### Core pathway signalling between cells: monodirectional or bidirectional?

While the involvement of Fz, Vang and Fmi in cell-cell signalling is well established, there has been considerable debate regarding whether information is transmitted between cells monodirectionally from Fz to Vang, or bidirectionally between Fz and Vang, with conflicting experimental data presented on each side.

Experiments in the *Drosophila* wing were designed to reveal the mechanism of signalling through examination of phenotypes around clones lacking both *Vang* and *fz* (Strutt and Strutt, 2007; Chen et al., 2008). It was hypothesised that should signalling be monodirectional from Fz-Fmi to Vang-Fmi, a double *Vang*^-^ *fz*^-^ clone would resemble a *fz*^-^ single clone. In this scenario, neighbouring cells would not ‘sense’ the lack of Vang within the clone and would only be affected by the lack of Fz activity (Fig.1E). However, if signalling were bidirectional, neighbouring cells would no longer be able to send or receive information to/from clonal cells that lack both *Vang* and *fz* and would thus have normal polarity (Fig.1F). In experiments, *Vang*^-^ *fz*^-^ clones showed little or no non-autonomy, suggesting a bidirectional mechanism (Strutt and Strutt, 2007; Chen et al., 2008).

However, a later study suggested that *Vang*^-^ *fz*^-^ clones both qualitatively and quantitatively gave the same phenotype as *fz*^-^ clones, supporting monodirectional signalling from Fz-Fmi to Fmi-Vang (Wu and Mlodzik, 2008). Taken together with biochemical data revealing a physical interaction between Fz and Vang, the authors concluded that Fz is a ligand for Vang, which acts as a receptor for polarising signals.

In studies on *Drosophila* abdomen hair polarity, it was also suggested that Fz-Fmi in one cell signals monodirectionally to Fmi-Vang in the next (Lawrence et al., 2004). However, on revisiting this work, Lawrence and colleagues concluded that an experimental artefact had misled them (Struhl et al., 2012) and a further series of experiments instead supported bidirectional signalling.

While the weight of evidence suggests bidirectional signalling is the likely mechanism, the conclusions drawn, particularly regarding experiments in the *Drosophila* wing, remain controversial. Furthermore, *fmi*^-^ single clones have been shown experimentally to show no non-autonomy in most cases and weak proximal non-autonomy in some examples (Chen et al., 2008; Le Garrec et al., 2006; Strutt and Strutt, 2007), but the mechanisms discussed here do not make predictions about the *fmi*^-^ clone phenotype within the context of mono-or bi-directional signalling.

### Mechanisms of feedback amplification of polarity

To generate a strongly polarised system, it is thought that small biases in protein localisation are induced by global cues (e.g. gradients), which are then amplified by positive feedback (Aw and Devenport, 2017). Such feedback is most commonly assumed to occur through ‘like’ complexes of the same orientation stabilising each other, and/or ‘unlike’ complexes of opposite orientations destabilising each other (Fig.1G). Both interactions are hypothesised to result in a local build-up of complexes of the same orientation (Klein and Mlodzik, 2005; Strutt and Strutt, 2009).

The non-transmembrane core pathway components, Dsh, Pk and Dgo, are required for amplification of polarity by promoting segregation of the core protein complexes to proximal and distal cell edges. Thus loss of their activity in clones of cells results in a failure of the mutant cells to planar polarise. However, this does not cause significant repolarisation of neighbouring wild-type cells (Gubb et al., 1999; Theisen et al., 1994; Amonlirdviman et al., 2005; Strutt and Strutt, 2007).

A number of molecular mechanisms have been proposed to mediate such stabilising and destabilising interactions. As core proteins progressively localise into clusters of the same orientation during polarisation, it has been suggested that ‘like’ complexes may intrinsically cluster (Strutt et al., 2011; Cho et al., 2015) and that this may be driven by multiple low-affinity interactions between core proteins leading to a phase transition into a stable state (Strutt et al., 2016). Conversely, destabilising interactions have been proposed to occur via Pk-Vang inhibiting Dsh binding to Fz (Tree et al., 2002; Amonlirdviman et al., 2005; Jenny et al., 2005), or by Pk reducing Dsh-Fz stability (Warrington et al., 2017), or by Fz-Dsh promoting Pk-mediated internalisation and turnover of Vang (Cho et al., 2015). Since such mechanisms suggest an effect on the stability of *inter*cellular complexes this might predict a role in altering signalling between cells, although the relationship between feedback and signalling directionality has not previously been examined.

### Computational modelling of planar polarity

Numerous computational models have been proposed, implementing a variety of different feedback interactions between core protein complexes, all of which successfully recapitulate a polarised state (e.g. Amonlirdviman et al., 2005; Le Garrec et al., 2006; Burak and Shraiman, 2009; Schamberg et al., 2010). However, while these models have generally attempted to reproduce single clone phenotypes, none have examined double clones or mechanisms of cell-cell signalling directionality. Furthermore, while it appears evident that core protein asymmetric distributions driven by feedback amplification are intrinsically linked to cell-cell signalling and propagation of planar polarity, the relationship between these phenomena remains largely unexplored.

To address these issues, we have used computational modelling to explore different scenarios for core pathway function, based on different assumptions regarding protein behaviours. This, taken together with new experimental data to reassess the *Vang*^-^ *fz*^-^ clone phenotype, allows us to make strong predictions regarding likely molecular mechanisms of action and provides the basis for future experimental studies.

## Results and Discussion

### A computational modelling framework for investigating cell-cell signalling and feedback amplification in planar polarity

We developed a computational framework to model potential molecular mechanisms for planar polarity signalling. The framework represents a simplified system, to allow rapid testing of different signalling regimes and simulation of clone phenotypes.

Within this framework, Fz and Vang each represent both the transmembrane protein encoded by the corresponding gene, and the associated cytoplasmic proteins with which they interact and which are required for feedback amplification of protein asymmetry. Thus, ‘Fz’ represents Fz bound to Dsh and Dgo, and ‘Vang’ represents Vang associated with Pk. Fmi is allowed to bind homophilically in trans between neighbouring cells (indicated by ‘:’ in later text), and also to interact in cis with either Fz or Vang in the membranes of the same cell (indicated by ‘-’ in later text; Fig.S1A).

Planar polarity was simulated in a one-dimensional row of cells, each with two compartments representing proximal (left) and distal (right) sides of the cell membrane, respectively (Fig.S1B). The system was initialised with one arbitrary unit (AU) of each molecular species per compartment at the start of each simulation, except with a small imbalance in initial Fz level towards distal sides of cells to provide a global orienting cue, which could be amplified by feedback interactions. Wild-type polarity was defined as higher Fz bound into complexes at distal cell ends and higher Vang bound at proximal cell ends (Fig.S1C).

As discussed, there is evidence for two forms of intracellular feedback interactions: local destabilisation of ‘unlike’ oriented complexes, and local stabilisation of ‘like’ oriented complexes (Fig.1G). Destabilising feedback interactions were implemented such that Vang-containing complexes (Vang-Fmi:Fmi, Vang-Fmi:Fmi-Fz and Vang-Fmi:Fmi-Vang) in each cell compartment destabilised Fz-containing complexes (Fz-Fmi:Fmi, Fz-Fmi:Fmi-Vang, Fz-Fmi:Fmi-Fz) in the same compartment, and vice versa. The strengths of destabilising feedbacks from Fz and Vang were given by the parameters *V*_max,F_ and *V*_max,V_, respectively, representing the maximum fold-change conferred to the off-rate of each reaction (see Experimental Procedures and Supplemental Text). Stabilising feedback interactions were implemented in a similar manner, by modulating reaction on-rates, with Vang- or Fz-containing complexes stabilising themselves.

After exploring both destabilising and stabilising feedback interactions in simulations, we concluded that in general these mechanisms polarised the system equivalently. However, while stabilising feedbacks recapitulated key clone phenotypes, there were subtle differences in some cases (e.g. Fig.S1D; Supplemental Text for further discussion). In particular, for systems relying only on stabilising feedbacks, ‘unlike’ complex stability was unchanged, thus the system was slower to polarise and more sensitive to the rate of protein diffusion to sort complexes. Hereafter, we describe models with destabilising feedbacks only. Since molecular evidence for local destabilising feedback interactions both from Fz to Vang and from Vang to Fz have been reported in the literature (Tree et al., 2002; Jenny et al., 2005; Cho et al., 2015; Warrington et al., 2017), both were implemented in these models, unless otherwise stated, by setting the values of *V*_max,F_ and *V*_max,V_ to greater than 1 (see Experimental Procedures; Fig.1H).

A set of ordinary differential equations describing the binding and diffusion events that occur in complex formation and localisation (Fig.S1B), were numerically solved and allowed to evolve to steady state (Supplemental Text).

We designed models with different signalling assumptions, namely ‘no direct signalling’ (Model 1), ‘direct monodirectional signalling’ (Model 2) or ‘direct bidirectional signalling’ (Model 3). While the molecular nature of such cell-cell signalling remains unclear, we have assumed that polarity information is transmitted directly via mass-action binding of complexes at intercellular junctions. To implement this we introduced variations in the relative dissociation constants of complexes, indicating how a cell may be able to directly ‘sense’ the presence of Fz or Vang in its neighbours when complexes form. For example in the no direct signalling model, both Fz and Vang were allowed to bind to and stabilise Fmi:Fmi dimers, such that all complexes involving Fz or Vang had equal K_D_ (Fig.2A). Thus, Fz and Vang could not promote each other’s incorporation into intercellular complexes and therefore could not ‘send’ information to neighbouring cells.

**Figure 2.**
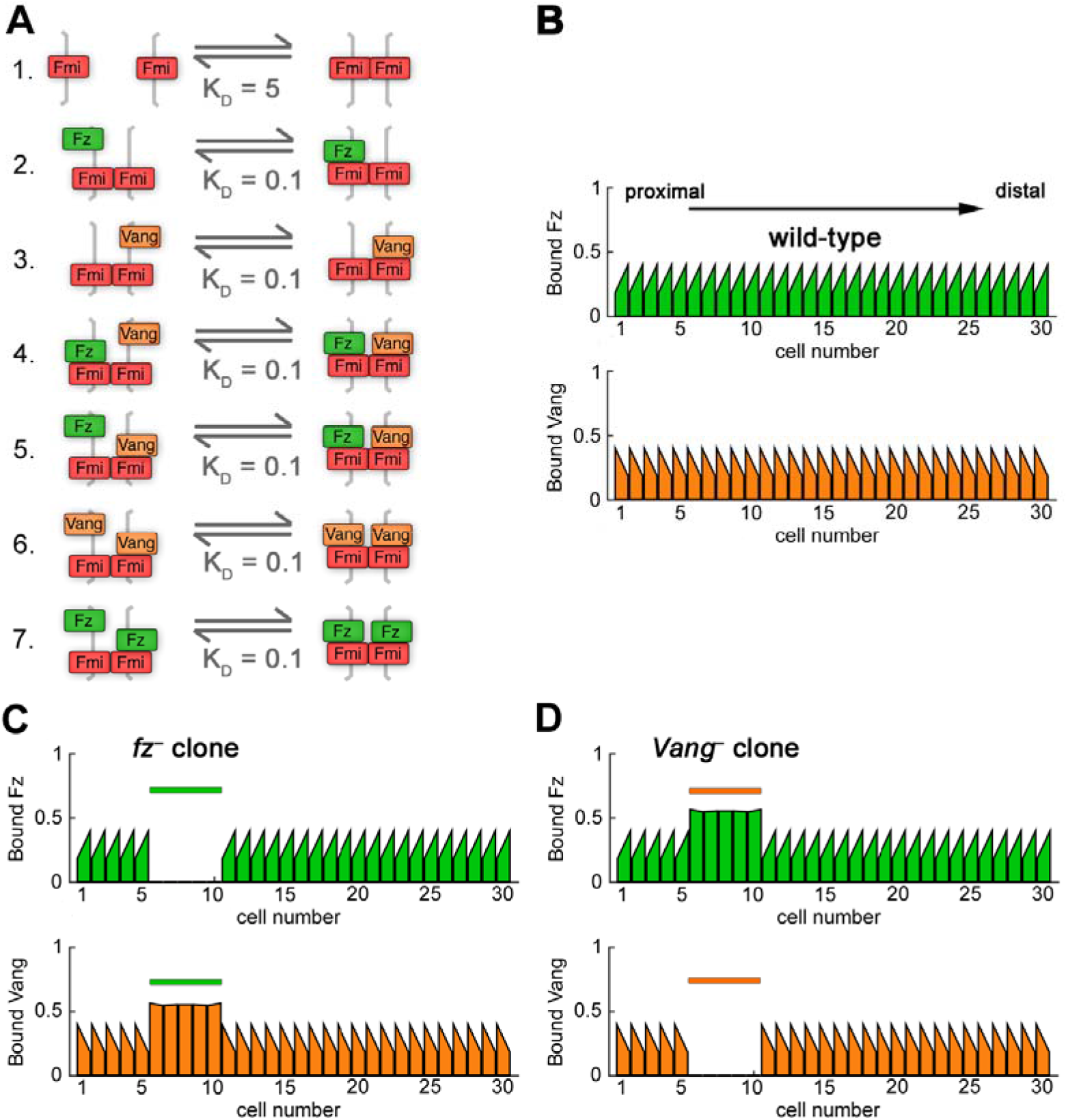
Polarity is generated in simulations with no direct signalling. (A) Model 1 biochemical binding reactions with relative dissociation constants K_D_. Higher K_D_ corresponds to weaker binding. (B) Simulation of wild-type polarity at steady state showing amount of bound Fz (top) and bound Vang (bottom) on the proximal and distal sides of each cell. Bound Fz levels (i.e. sum of complexes that contain Fz within each compartment; upper panels, green graphs) and bound Vang levels (lower panels, orange graphs) are shown for each cell edge. (C,D) Simulation at steady state (*V*_max,F_ = *V*_max,V_ =10) for *fz*^-^ (C) or *Vang*^-^ (D) clones. Sloped tops of bars indicate the cell is polarised for that protein. Coloured bars above graphs indicate clone cells (in cell numbers 6-10). Cells neighbouring the clones show normal polarity and thus clones are autonomous.

We tested each model’s ability to reproduce the following experimental observations:

i. polarisation of Fz and Vang in wild-type tissue;
ii. reversal of polarity in the 5-10 cells neighbouring those in a clone lacking the activity of *fz* or *Vang*;
iii. no reversal of polarity in cells neighbouring a *fmi*^-^ clone.

Models that met each of these criteria were then used to predict the phenotype of *Vang*^-^ *fz*^-^ double clones, and these predictions were then tested experimentally. We quantified non-autonomy around a clone as the number of cells with reversed polarisation in terms of Fz localisation (in general results were the same if we instead considered Vang localisation [e.g. Fig.3B]). Reversals of polarity in cells to the right of a clone, such that Fz pointed towards the clone, was termed distal non-autonomy (i.e. a *fz*-like phenotype), while polarity reversals to the left were termed proximal non-autonomy (*Vang*-like).

**Figure 3.**
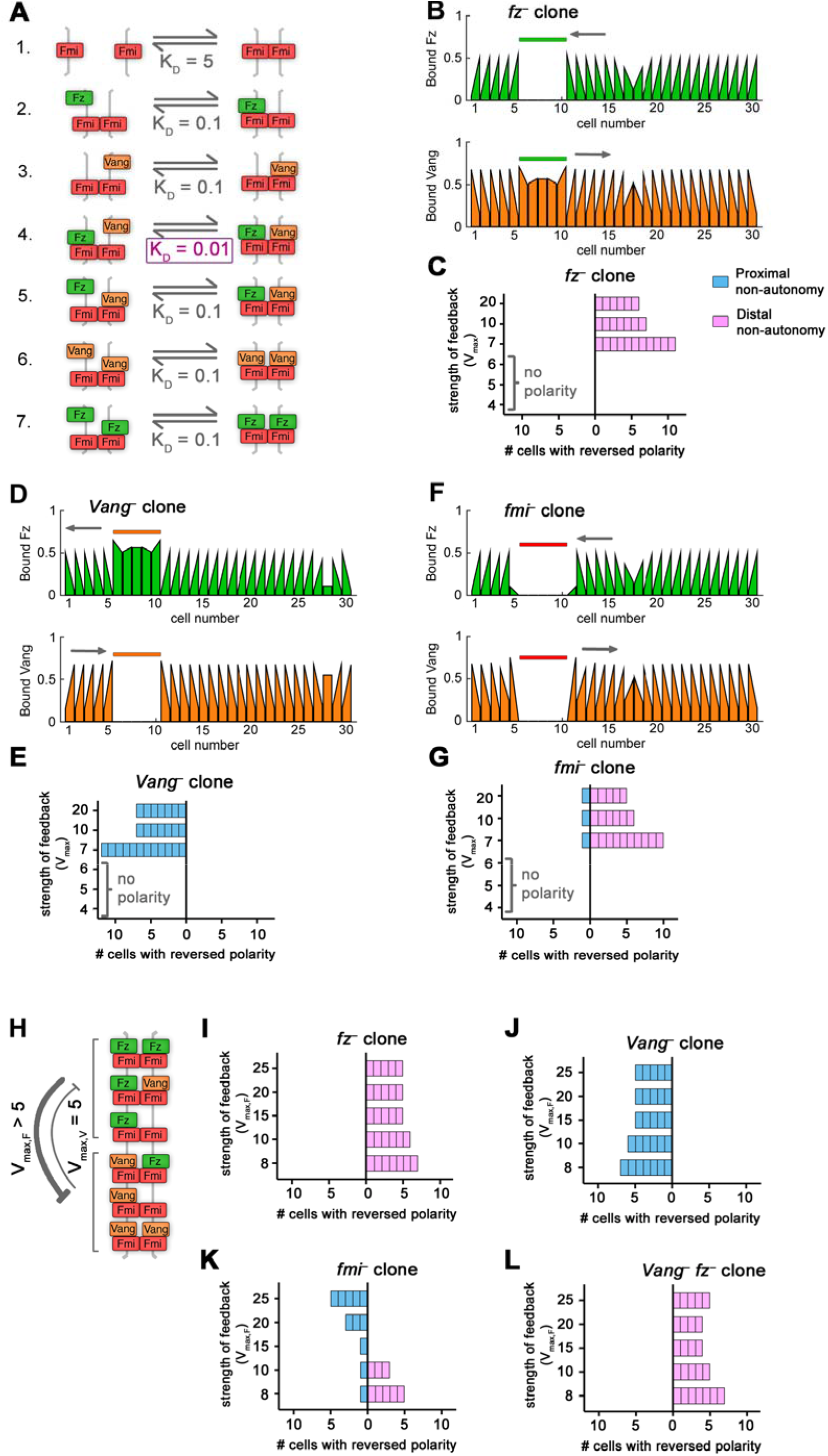
Direct monodirectional signalling can reproduce *in vivo* clone phenotypes. (A) Model 2 biochemical binding reactions with relative dissociation constants, K_D_, for direct monodirectional signalling. In this model, Vang is better at binding (i.e. has a lower dissociation constant) to complexes that already have Fz bound in the neighbouring cell (reaction 4) compared to other complexes (reactions 3 and 6). This allows Vang to receive a ‘signal’ from Fz. (B,D,F) Bound Fz levels (upper panels, green graphs) and bound Vang levels (lower panels, orange graphs) at each cell edge from simulations, with equal feedback strengths (*V*_max,F_ = *V*_max,V_ = 10), at steady state for *fz*^-^ (B), *Vang*^-^ (D) or *fmi*^-^ (F) clones. Sloped tops of bars indicate the cell is polarised for that protein. Coloured bars above graphs indicate clone cells (in cell numbers 6-10). Grey arrows indicate regions of non-autonomous polarity. (C,E,G) Non-autonomy around clones from simulations at steady state with varying, but balanced, feedback strength (where *V*_max_ represents *V*_max,F_ *= V*_max,V_). Results shown for *fz*^-^ (C), *Vang*^-^ (E), or *fmi*^-^ (G) clones. Parameter conditions where no polarity was observed in the absence of clones are indicated (‘no polarity’). (H) Diagram indicating unbalanced feedback interaction strengths with stronger feedback from Fz as compared to Vang (*V*_max,F_ > *V*_max,V_). (I-L) Simulations of monodirectional signalling with unbalanced feedback strengths as in (H), where *V*_max,V_ = 5 and *V*_max,F_ > 5, show distal and proximal non-autonomy around *fz*^-^ (I) and *Vang*^-^ (J) clones, respectively. However, *fmi*^-^ clones (K) show non-autonomy varying from distal to proximal as *V*_max,F_ increases, while *Vang*^-^ *fz*^-^ double clones (L) show distal non-autonomy for all parameters shown. If *V*_max,V_ = 5, *fmi*^-^ clones show no non-autonomy only in the case *V*_max,F_ = 15; in this case, double clones show distal non-autonomy.

### A model with no direct signalling is unable to reproduce non-autonomy around *fz*^-^ and *Vang*^-^ clones (Model 1)

We first assessed whether direct signalling is required to establish polarity in our modelling framework by simulating complex formation with no direct communication between cells. We termed this model ‘no direct signalling’ (Table 1: Model 1; Fig.2), indicating that a cell is unable to directly ‘sense’ the presence of Fz or Vang in its neighbours when complexes form. As discussed previously, this is achieved by implementing equal K_D_ for complex formation in reactions 2-7 (Fig.2A). Thus, Fz and Vang do not directly promote each other’s incorporation into intercellular complexes.

**Table 1.**
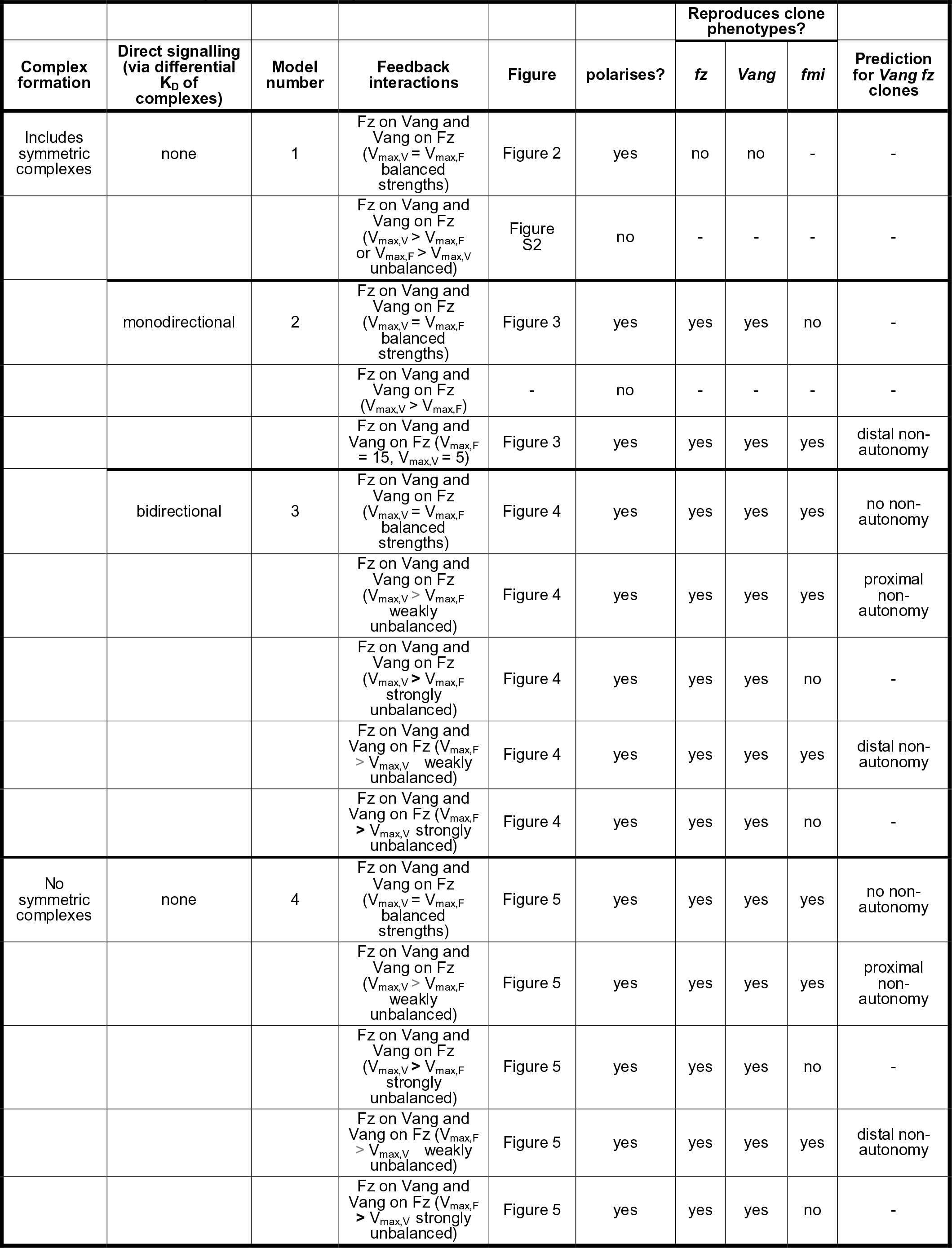
Summary of planar polarity models.

We established parameter values (see Supplemental Text) for which Model 1 generated wild-type polarity (Fig.2B), thus meeting criterion (i). We then examined the behaviour of loss-of-function clones in this model. We hypothesised that under the ‘no signalling’ assumption, this model would not generate non-autonomy around clones of cells with altered Fz or Vang activity.

Indeed, we found no non-autonomy around *fz*^-^ or *Vang*^-^ clones (Fig.2C,D), thus this model failed to meet criterion (ii). Increasing the strength of feedback did not result in non-autonomy around such clones, while decreasing it below a threshold resulted in no polarisation. To rationalise this, we consider the possible complexes that can form on the boundary of a *fz*^-^ clone. Allowing all complexes to form with equal affinity leads to equal amounts of Fz and Vang on the outer clone boundary and thus no preferential accumulation of either protein over the other (Fig.S2A). In this model, since Fz or Vang binding in one cell is not influenced by availability of Fz or Vang in a neighbouring cell, in cells outside the clone the initial bias in unbound Fz levels is the driving force for feedback-amplified polarisation.

We next relaxed our simplifying assumption that feedback interactions act with equal strength and examined the model with unbalanced feedbacks (*V*_max,F_ ≠ *V*_max,V_) or with just a single feedback (where either *V*_max,F_ or *V*_max,V_ are 1). For the initial Fz bias considered, simulations revealed that the model failed to polarise if feedbacks were unbalanced (e.g. Fig.S2B,C). Based on its inability to reproduce observed *fz*^-^ and *Vang*^-^ clone phenotypes, we rejected Model 1, and proceeded to consider the effect of direct signalling.

### A direct monodirectional signalling model reproduces non-autonomous phenotypes around *fz*^-^, *Vang*^-^ and *fmi*^-^ clones and predicts distal non-autonomy around *Vang*^-^ *fz*^-^ double clones (Model 2)

We next implemented ‘direct monodirectional signalling’ (Table 1: Model 2; Fig.3), from Fz to Vang, such that Vang bound more strongly to a Fmi:Fmi-Fz complex (i.e. a low dissociation constant; Fig.3A, reaction 4) than to just Fmi:Fmi (Fig.3A, reaction 3), thereby ‘sensing’ the presence of Fz in the neighbouring cell. In contrast, Fz bound to a Fmi:Fmi-Vang complex (Fig.3A, reaction 5) had the same intermediate dissociation constant as Fz bound to just Fmi:Fmi (Fig.3A, reaction 2). Thus localisation of Vang at junctions was promoted by Fz in the next cell, but Fz was unaffected by Vang in the next cell (Fig.3A). As for Model 1, we considered polarity to be amplified by mutually destabilising feedback interactions, initially acting with equal strength (see Fig.1H).

Monodirectional signalling resulted in polarisation of wild-type cells and furthermore, reproduced the experimentally observed distal and proximal non-autonomy around *fz*^-^ and *Vang*^-^ clones, respectively, for a range of feedback strengths (Fig.3B-E), thus meeting criteria (i) and (ii). We rationalise this by considering that Vang binds preferentially to complexes containing Fz in the next cell. Thus, in cells immediately neighbouring a *fz*^-^ clone, Vang preferentially localises away from the clone (Fig.S3A) which results in reversed polarity distal to the clone.

The range of non-autonomy was a function of the strength of feedbacks (*V*_max_, where *V*_max_ = *V*_max,F_ = *V*_max,V_). Higher values of either parameter suppressed non-autonomy, since the amplification of the initial bias in each cell dominated over the mislocalisation of protein complexes propagating from the clone edge (Fig.3C,E; see Supplemental Text).

Simulations revealed that our model of direct monodirectional signalling showed distal non-autonomy around *fmi*^-^ clones for a range of feedback strengths (Fig.3F,G) and therefore did not recapitulate experimental observations in the fly wing. We note that in the first cell neighbouring the clone, neither Vang nor Fz could bind at the clone boundary (Fig.3F). Thus, since we have quantified non-autonomy via Fz localisation, the first cell proximal to the clone is also scored as non-autonomous (Fig.3G).

To rationalise the distal non-autonomy, we consider complexes that can form at a *fmi*^-^ clone boundary (Fig.S3B). In cells immediately next to the clone, both Fz and Vang localise away from the clone. However, Vang preferentially binds to Fz-containing complexes and thus accumulates on the boundary furthest from the clone to higher levels than Fz and this difference is amplified by the feedback interactions (Fig.S3B, orange arrows), leading to distal non-autonomy. The failure to mimic experimental observations led us to reject Model 2 under the conditions of balanced feedbacks.

We next relaxed our simplifying assumption that feedback interactions act with equal strength and examined the model with unbalanced feedbacks (*V*_max,F_ ≠ *V*_max,V_) or with just a single feedback (where either *V*_max,F_ or *V*_max,V_ are 1). Simulations revealed that this model could no longer generate a polarised steady state when feedback was stronger from Vang (*V*_max,F_ < *V*_max,V_), or with either feedback operating alone, thereby failing to meet criterion (i). However, with stronger feedback acting from Fz complexes to destabilise Vang binding (V_max,F_ > V_max,V_, where *V*_max,V_ ≥ 5; e.g. Fig.3H), this model generated a polarised steady state and recapitulated the phenotypes of *fz*^-^ and *Vang*^-^ (Fig.3I,J). For a limited parameter range (*V*_max,F_ = 15, *V*_max,V_ = 5), this model also generated autonomous *fmi*^-^ clones (Fig.3K), thereby meeting all of our criteria.

Having found conditions (i.e. with stronger feedback from Fz) under which the direct monodirectional signalling model could reproduce criteria (i)-(iii), we then used it to predict the phenotype of *Vang*^-^ *fz*^-^ double clones, for which experimental results remain controversial (Strutt and Strutt, 2007; Chen et al., 2008; Wu and Mlodzik, 2008). Simulations revealed distal non-autonomy around such double clones (Fig.3L). In cells immediately neighbouring *Vang*^-^ *fz*^-^ clones, Vang preferentially binds to Fz-containing complexes and thus accumulates on the boundary furthest from the clone (Fig.S3C, orange arrows). Based on these simulated clone phenotypes, we conclude that Model 2 may be a valid model of planar polarisation in the fly wing if feedback is stronger from the Fz side of the complex, and it predicts distal non-autonomy around *Vang*^-^ *fz*^-^ double clones.

### A direct bidirectional signalling model reproduces non-autonomous phenotypes around *fz*^-^, *Vang*^-^ and *fmi*^-^ clones and predicts no non-autonomous polarity around *Vang*^-^ *fz*^-^ double clones (Model 3)

An alternative mechanism of signalling that has been presented in the literature is that of bidirectional signalling. We next established a model of ‘direct bidirectional signalling’ (Table 1: Model 3; Fig.4) to address whether it too could meet our criteria of generating a polarised steady state and reproducing single clone phenotypes. In this model both Vang bound to a Fmi:Fmi-Fz complex (Fig.4A, reaction 4) and Fz bound to a Fmi:Fmi-Vang complex (Fig.4A, reaction 5) had low dissociation constants, but Vang bound to Fmi:Fmi (Fig.4A, reaction 3) and Fz bound to Fmi:Fmi (Fig.4A, reaction 2) had intermediate dissociation constants. Thus Fz and Vang both promoted each other‘s binding in the next cell (Fig.4A). As for our previous models, we considered amplification to be mediated by mutually destabilising feedback interactions, initially acting with equal strengths (see Fig.1H).

**Figure 4.**
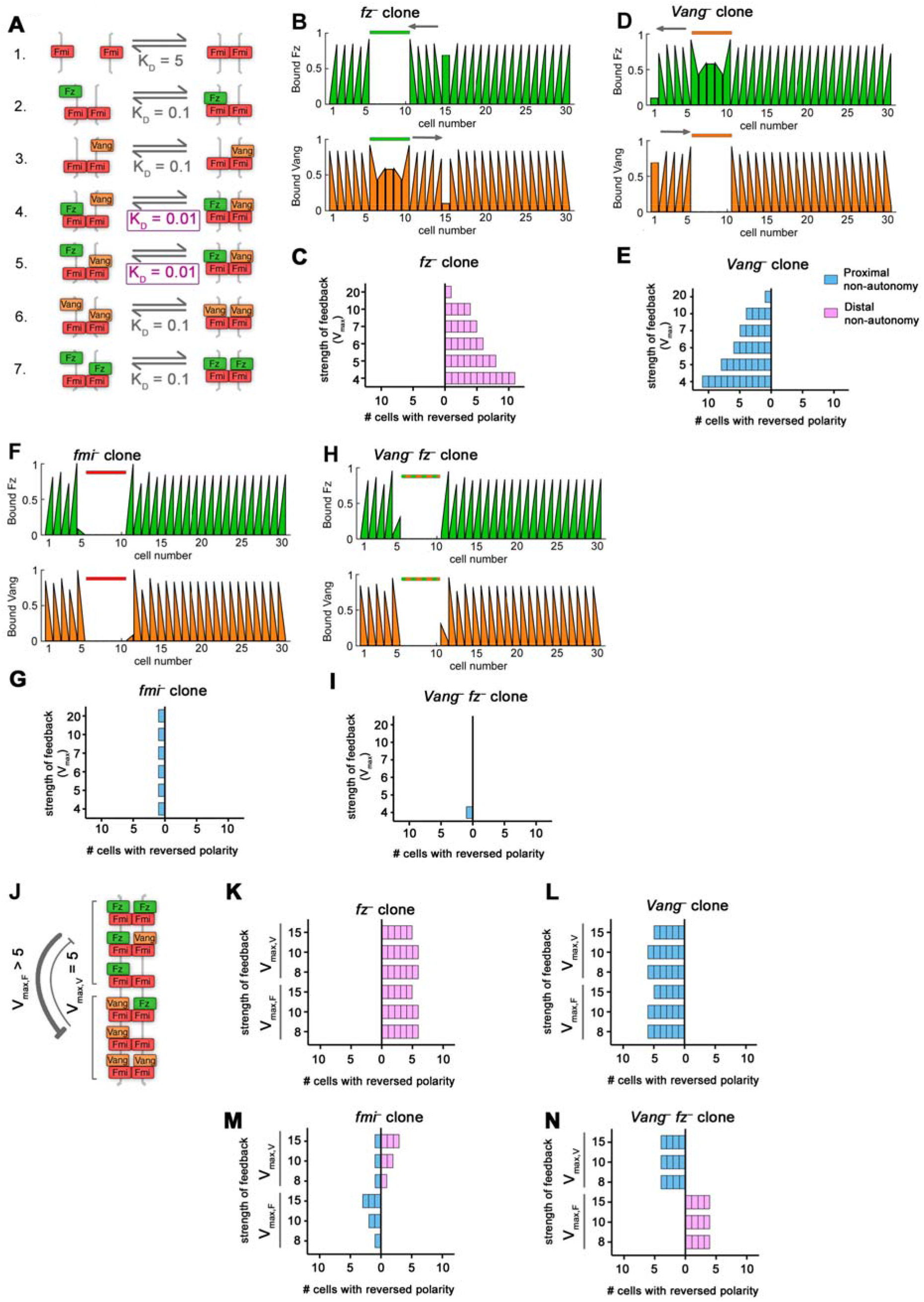
Direct bidirectional signalling reproduces *in vivo* clone phenotypes. (A) Model 3 biochemical binding reactions with relative Dissociation Constants, K_D_, for direct bidirectional signalling. In this model, Vang is better at binding (i.e. has a lower dissociation constant) to complexes that already have Fz bound in the neighbouring cell (reaction 4) compared to other complexes (reactions 3 and 6) and Fz is *also* better at binding when Vang is present in the complex (reaction 5 compared to 2 and 7). This allows both Vang and Fz to receive information in the form of mass action binding kinetics. (B,D,F,H) Bound Fz levels (upper panels, green graphs) and bound Vang levels (lower panels, orange graphs) at each cell edge from simulations at steady state for one parameter set (*V*_*max,F*_ = *V*_*max,V*_ = 10) for *fz*^-^ (B), *Vang*^-^ (D), *fmi*^-^ (F), or *Vang*^-^ *fz*^-^ (H) clones. Coloured bars above graphs indicate clone cells (in cell numbers 6-10). Grey arrows indicate regions of non-autonomous polarity. (C,E,G,I) Non-autonomy around clones from simulations at steady state with varying, but balanced, feedback strength (where *V*_max_ represents *V*_max,F_ = *V*_max,V_). Results shown for *fz*^-^ (C), *Vang*^-^ (E), *fmi*^-^ (G), or *Vang*^-^ *fz*^-^ (I) clones. (J) Diagram showing an example of unbalanced intracellular destabilising feedbacks between Fz and Vang. In this example there is stronger feedback from Fz than from Vang. (K-N) Non-autonomy around clones from simulations at steady state with unbalanced feedback strength for *fz*^-^ (K), *Vang*^-^ (L), *fmi*^-^ (M), or *Vang*^-^ *fz*^-^(N). For the top half of each graph *V*_max,V_ is >5 as indicated and *V*_max,F_ = 5. For the lower bars of each graph, *V*_max,F_ is > 5 as indicated and *V*_max,V_ = 5. Clones of *fz*^-^ and *Vang*^-^ show distal and proximal non-autonomy, respectively. However, *fmi*^-^ clones show distal non-autonomy when *V*_max,F_ is higher, but proximal non-autonomy when *V*_max,V_ is higher. The direction of non-autonomy is reversed for *Vang*^-^ *fz*^-^ double clones, showing distal non-autonomy when *V*_max,V_ is higher, but proximal non-autonomy when *V*_max,F_ is higher.

In this model the expected distal and proximal non-autonomy were observed around *fz*^-^ and *Vang*^-^ clones, respectively (Fig.4B-E). We rationalise these phenotypes by considering the complexes that can form in the cells immediately neighbouring the clone. For example, in cells next to a *fz*^-^ clone, Vang preferentially localises towards cells expressing its binding partner, Fz. Since there is no Fz within the clone, Vang accumulates on boundaries furthest from the clone (Fig.S4A, orange arrows). This model also recapitulated the expected phenotype of *fmi*^-^ clones, for which neither Fz nor Vang binding is favoured at clone boundaries, resulting in no propagating non-autonomy (Fig.4F,G; Fig.S4B).

Since the direct bidirectional signalling model accurately reproduced *fz*^-^, *Vang*^-^ and *fmi*^-^ clone phenotypes, meeting criteria (i)-(iii), we used this model to predict the phenotype of *Vang*^-^ *fz*^-^ double clones. In contrast to the direct monodirectional model, it predicted no non-autonomous polarity around *Vang*^-^ *fz*^-^ double clones (Fig.4H,I). Considering the complexes that can form at this clone boundary, both Fz and Vang can form trimer complexes on the edge of the clone, but these are less favoured than tetramer complexes. Thus both Fz and Vang preferentially localise to the boundary furthest from the clone where neither is favoured over the other (Fig.S4C). Polarity direction is thus driven by the initial bias in Fz localisation, not by the clone.

To examine whether our findings depended on our simplifying assumption that feedback interactions act with equal strength, we relaxed this assumption and examined the model with unbalanced feedbacks (*V*_max,F_ ≠ *V*_max,V_; for example Fig.4J) or with just a single feedback (where either *V*_max,F_ or *V*_max,V_ are 1). If only a single feedback was present, the system did not polarise. In the case of unbalanced feedbacks, single *fz*^-^ or *Vang*^-^ clones behaved as expected, with distal and proximal non-autonomy (Fig.4K,L), respectively.

For strong differences in feedback strength (i.e. *V*_max,F_ = 5, *V*_max,V_ > 8 or *V*_max,V_ = 5, *V*_max,F_ > 8), *fmi*^-^ clones exhibited non-autonomy (Fig.4M), thus failing to meet criterion (iii). In cells neighbouring *fmi*^-^ clones, the immediate boundary is unable to localise any complexes due to the inability to form Fmi:Fmi dimers. Since all of the proteins in such cells must localise to the boundary furthest from the clone, the protein mediating the feedback ‘wins’ on this boundary (Fig.S5A). For instance, Fz accumulates on the boundary furthest from the clone when it more strongly destabilises Vang, and Vang accumulates away from the clone when it strongly destabilises Fz. In contrast, for weak differences in feedback strength (e.g. *V*_max,F_ = 5, *V*_max,V_ = 8) little non-autonomy was observed in cells neighbouring *fmi*^-^ clones (Fig.4M), thus meeting criterion (iii). We therefore examined the phenotype predicted for *Vang*^-^ *fz*^-^ double clones under these conditions. They showed distal non-autonomy when feedback was slightly stronger from Fz, but proximal non-autonomy when feedback was slightly stronger from Vang (Fig.4N; Fig.S5B).

Based on these simulated clone phenotypes, we conclude that Model 3 may be a valid model of planar polarisation in the fly wing. If feedback is balanced between the two sides of the complex, criteria (i)-(iii) are met and no non-autonomy is predicted around *Vang*^-^ *fz*^-^ double clones. However, all criteria are also met if feedbacks are weakly unbalanced, but this leads to distal or proximal non-autonomy around *Vang*^-^ *fz*^-^ double clones depending on which feedback is strongest.

### The absence of symmetric complex formation leads to indirect signalling (Model 4)

Notably there is no experimental evidence to rule out the formation of symmetric complexes (Fz-Fmi:Fmi-Fz and Vang-Fmi:Fmi-Vang), and visualising the structure of individual complexes is beyond the limits of conventional microscopy. Therefore, we allowed symmetric complexes to form in Models 1-3. However, previously published computational models have assumed that symmetric complexes do not form (Amonlirdviman et al., 2005; Le Garrec et al., 2006; Burak and Shraiman, 2009). To test whether this difference was important, we adapted our model with no direct signalling (Model 1), to block the formation of symmetric complexes (Table 1: Model 4; Fig.5). This was achieved by setting the relevant binding rate constants (*k*_6_, *k*_7_) to zero (Fig.5A). We first confirmed that this model could generate stably polarised cells through amplification of the initial global cue by local destabilising feedback interactions both from Fz to Vang and from Vang to Fz (Fig.5B), thereby meeting criterion (i).

**Figure 5.**
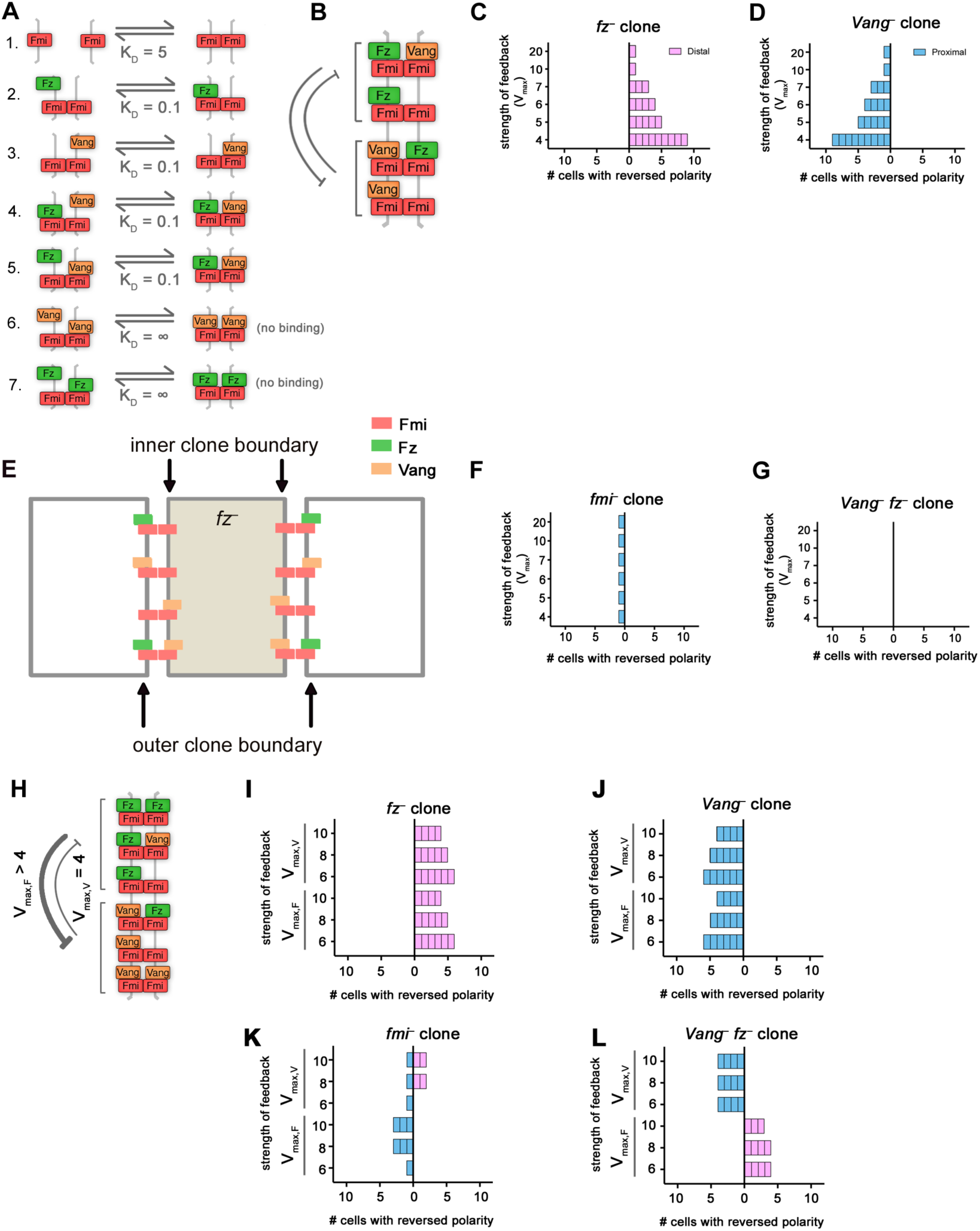
In the absence of symmetric complexes feedback interactions determine directionality of indirect signalling. (A) Model 4 biochemical binding reactions with relative dissociation constants K_D_. Symmetric complexes in reactions 6 and 7 do not form. (B) Diagram showing bidirectional intracellular destabilising feedbacks between Fz and Vang implemented in this model. (C,D,F,G) Non-autonomy around clones from simulations at steady state for *fz*^-^ (C), *Vang*^-^ (D), *fmi*^-^ (F), or *Vang*^-^ *fz*^-^ (G). Feedback strength varies as indicated (where *V*_max_ = *V*_max,F_ = *V*_max,V_). (E) Diagram of complex formation with ‘no direct signalling’ in the absence of symmetric complexes at the boundary of a *fz*^-^ clone. There are more possibilities for Fz than Vang to bind at the outer clone boundaries, which generates distal non-autonomy. (H) Diagram showing an example of unbalanced intracellular destabilising feedbacks between Fz and Vang. In this case there is stronger feedback from Fz than from Vang. (I-L) Non-autonomy around clones from simulations at steady state with unbalanced feedback strength for *fz*^-^ (I), *Vang*^-^ (J), *fmi*^-^ (K), or *Vang*^-^ *fz*^-^ (L). For the top half of each graph *V*_*max*,V_ is indicated as >4 and *V*_*max*,F_ = 4. For the lower bars of each graph, *V*_*max*,F_ is indicated as >4 and *V*_*max*,V_ = 4.

We next introduced clones into the model. In agreement with previous modelling work and despite the lack of direct signalling across complexes, we found that *fz*^-^ clones showed distal non-autonomy (Fig.5C) and *Vang*^-^ clones showed proximal non-autonomy (Fig.5D), thereby meeting criterion (ii). To illustrate the consequences of only allowing asymmetric complexes to form, we consider the example of a *fz*^-^ clone. On the boundary between a *fz*^-^ clone and a neighbouring wild-type cell, only Vang can bind to Fmi:Fmi on the inner clone boundary. At the outer clone boundary Fz binding is favoured over Vang binding (as Vang-Fmi-Fmi-Vang complexes could not form, Fig.5E). This generates an imbalance whereby more Fz can bind to the outer clone boundary than Vang and this difference can be amplified by feedback interactions (Fig.5E).

Simulations revealed that *fmi*^-^ clones were autonomous for this model (Fig.5F). Since this model was able to reproduce all single clone phenotypes meeting criteria (i)-(iii), we went on to use it to predict the outcome of *Vang*^-^ *fz*^-^ clones. Such clones were autonomous (Fig.5G), mimicking the findings of the direct bidirectional signalling mode (Model 3).

We next relaxed our assumption that feedbacks operate with equal strengths and examined the model with unbalanced feedbacks (*V*_max,F_ ≠ *V*_max,V_; for example Fig.5H) or with just a single feedback (where either *V*_max,F_ or *V*_max,V_ are 1). If only a single feedback was present, the system did not polarise. We found that single *fz*^-^ or *Vang*^-^ clones behaved as expected when feedbacks were unbalanced (Fig.5I,J).

As for Model 3, weak differences in feedback strength (e.g. *V*_max,F_ = 4, *V*_max,V_ = 6 or *V*_max,F_ = 6, *V*_max,V_ = 4) resulted in little non-autonomy in cells neighbouring *fmi*^-^ clones (Fig.5K), and criteria (i)-(iii) were all met. Thus, we examined the phenotype predicted for *Vang*^-^ *fz*^-^ double clones under these conditions, finding them to show proximal non-autonomy when feedback from Vang was stronger, but distal non-autonomy when feedback from Fz was stronger (Fig.5L). For stronger differences in feedback strength, *fmi*^-^ clones exhibited non-autonomy (Fig.5K), thus failing to meet criterion (iii).

We conclude that Model 4 may be a valid model of planar polarity in the fly wing and that the presence of two opposing feedback interactions results in ‘indirect bidirectional signalling’ (with the balance of the feedbacks determining the balance of the directionality).

To summarise our findings so far, we can find regions in parameter space for the direct monodirectional (Model 2), direct bidirectional (Model 3) and indirect bidirectional (Model 4) models that recapitulate *fz*^-^, *Vang*^-^ and *fmi*^-^ single clone phenotypes. However, these models predict qualitative differences in the phenotype around *Vang*^-^ *fz*^-^ double clones. To constrain our models and identify the most likely mechanism of planar polarity signalling in the wing, we re-examined the *Vang*^-^ *fz*^-^ double clone phenotype.

### The non-autonomous phenotype of *fz*^-^ clones is suppressed by simultaneous loss of *Vang*

As the *fz* and *Vang* genes lie on different chromosomes, previous studies generated clones of double-mutant tissue using transgenes that artificially provided *fz* or *Vang* function on a different chromosome arm. These studies used exogenous promoters, which might not provide identical activity levels to the endogenous genes (Strutt and Strutt, 2007; Chen et al., 2008; Wu and Mlodzik, 2008). Thus in each study, cells lacking Fz and Vang activity are juxtaposed to neighbours with potentially differing levels of Fz and Vang activity, and this may explain the varying degrees of non-autonomous propagation of polarity reported in each case.

To circumvent the disadvantages inherent in this approach, we generated *fz*^-^ clones in which Vang activity was either normal or was reduced only within the cells of the clone using RNAi in a MARCM GAL4/GAL80-dependent strategy (Lee and Luo, 1999 see Experimental Procedures). In this method, GAL80 suppresses expression of the RNAi transgene in all tissue except for the clone tissue. No transgenes were used to substitute for Fz or Vang activity, and thus the different clone genotypes should be directly comparable.

Control *fz*^-^ clones in the pupal wing, examined shortly before or around the time of trichome formation, showed the expected strong Fz and Vang localisation at clone boundaries (Fig.6A, white arrowheads). Propagation of this aberrant polarity extended around 5-10 cells into neighbouring tissue with associated mispolarisation of trichomes pointing towards clone tissue (Fig.6B, arrows, Vinson and Adler, 1987; Strutt, 2001; Bastock et al., 2003).

**Figure 6.**
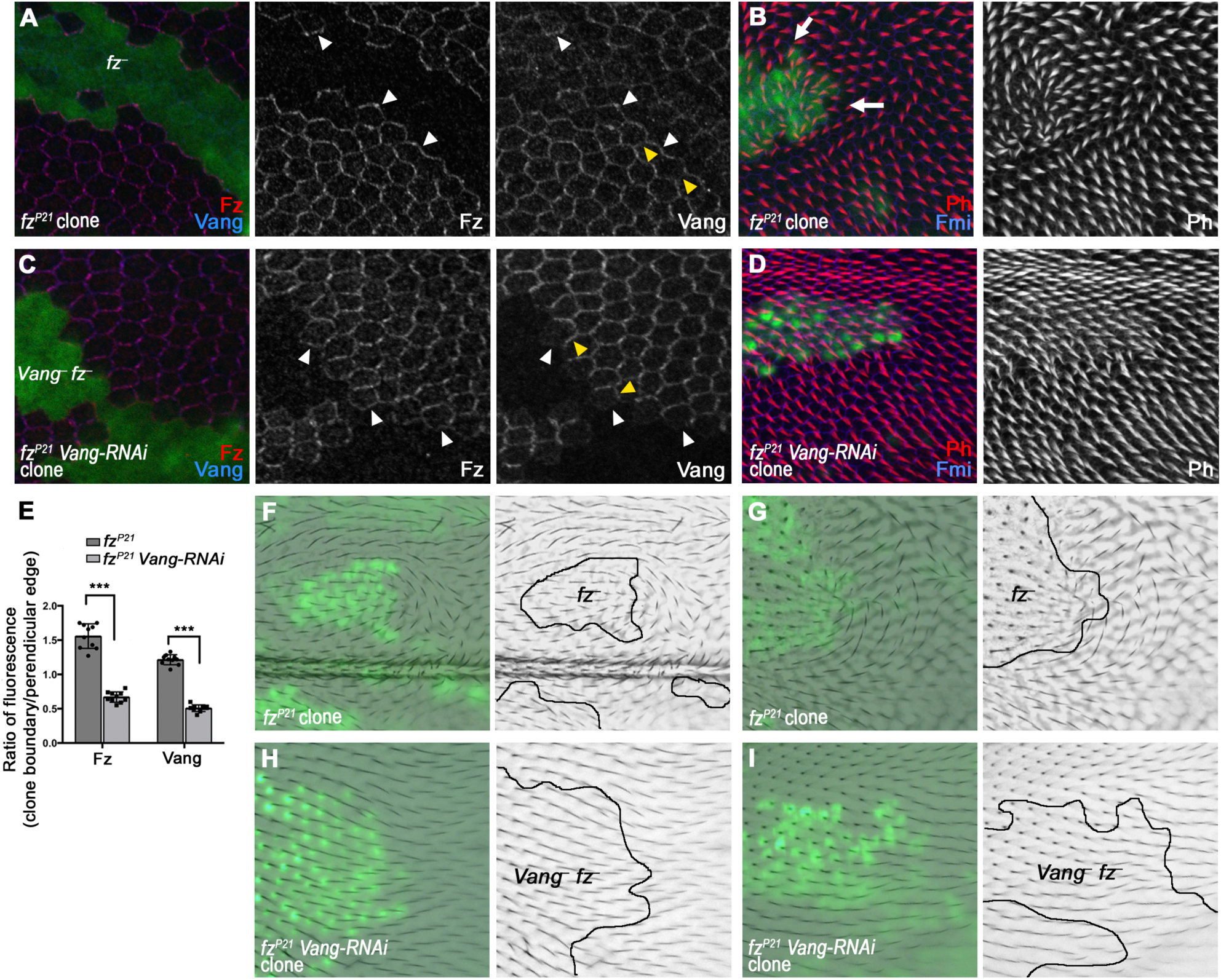
Non-autonomy around *fz*^-^ clones is suppressed by simultaneous knockdown of *Vang* activity by RNAi. (A-D) Pupal wings with loss-of-function MARCM clones of *fz*^*P21*^ alone (A,B), or *fz*^*P21*^ clones also expressing *UAS-Vang-RNAi*. Wings were immunolabelled for Fz (red) and Vang (blue) at 28 h after pupa formation (APF; A,C), or for Phalloidin to mark trichomes (red) and Fmi (blue) at 32.25 h APF (B,D). White arrowheads point to clone boundaries and yellow arrowheads point to cell edges perpendicular to the clone boundary. White arrows show areas of reversed trichome polarity. GFP (green) positively marks the clones. Representative images from at least 9 clones per genotype are shown. (E) Ratio of Fz and Vang fluorescence intensities at clone boundaries (white arrowheads in A and C) and perpendicular edges (yellow arrowheads in A and C). For *fz*^-^ clones (dark grey bars), Fz and Vang are recruited to the clone boundaries and so ratio values are above 1. However, for *Vang*^-^ *fz*^-^ clones (light grey bars), Fz and Vang do not accumulate at the boundary and thus ratio values are below 1. Measurements were taken from 10 wings per genotype (n = 10), averaged from at least 7 regions of each type per wing. Two-way ANOVA was employed with Sidak’s multiple comparison test, *** *p* < 0.0001. (F-I) Adult wings with clones of *fz*^*P21*^ (F,G) or *fz*^*P21*^ *UAS-Vang-RNAi* (H,I). Green nuclei mark clone cells and brightfield images show trichomes. Two representative examples from at least 10 clones are shown for each genotype. All panels are aligned with proximal left and anterior up.

In *fz*^-^ clones, expressing a *UAS-Vang-RNAi* transgene caused loss of immunoactivity for Vang protein (Fig.6C). There was no notable effect on trichome polarity outside the clones (Fig.6D) and consistent with this, Fz and Vang recruitment to the clone boundary (Fig.6C, white arrowheads) from other cell edges (Fig.6C, yellow arrowheads) was suppressed (Fig.6E).

We further considered whether the differences seen in previous reports were due to the stage at which trichome polarity was assayed, as two reports looked at pupal stages (Strutt and Strutt, 2007; Chen et al., 2008) and one analysed adult wings (Wu and Mlodzik, 2008). However, looking in adult wings in which Vang activity in *fz*^-^ clones was reduced by RNAi, we again observed a strong suppression of non-autonomy (compare Fig.6F,G to Fig.6H,I).

Thus, reduction in Vang activity in *fz*^-^ clones reduces Fz and Vang localisation at clone boundaries, and reduces propagation of polarity defects outside the clone, supporting previous reports that the non-autonomy of *Vang*^-^ *fz*^-^ clones is suppressed as compared to *fz*^-^ clones (Strutt and Strutt, 2007; Chen et al., 2008). Together with our modelling predictions this suggests that signalling is bidirectional. Thus we reject Model 2 and conclude that Models 3 and 4 are plausible models of planar polarity signalling in the wing.

## Conclusions

We set out to understand the possible molecular wirings that might underlie the cell-cell coordination of planar polarity. We specifically address possible scenarios that would explain the observed behaviour of the core planar polarity pathway in the *Drosophila* wing, but our findings also reveal other scenarios that could coordinate polarity in other contexts.

We developed a suite of models reflecting different ‘signalling’ regimes that could reproduce a wild-type polarised steady state and single clone phenotypes (Table 1). Our monodirectional and bidirectional signalling models were able to produce polarised tissue and reproduce the phenotypes of single mutant *fz*^-^ and *Vang*^-^ clones. Interestingly, monodirectional and bidirectional models made different predictions about the feedback interactions necessary to reproduce the autonomy of *fmi*^-^ clones. Namely, monodirectional signalling from Fz to Vang required that feedback be stronger from Fz than Vang, while bidirectional signalling required that feedbacks be balanced (or only weakly unbalanced).

Under such conditions, monodirectional and bidirectional models made different predictions about the *Vang*^-^ *fz*^-^ double clone phenotype, either predicting distal non-autonomy or complete autonomy. By carrying out new experiments examining the double *Vang*^-^ *fz*^-^ clone phenotype in the wing, we confirm that non-autonomy is suppressed when compared to either *fz*^-^ or *Vang*^-^ single mutant clones. This supports the conclusion that signalling between Fz and Vang is bidirectional and that feedback interactions are balanced, with Models 3 and 4 thus being the most plausible.

Interestingly, simulations revealed that bidirectional signalling could be mediated either *indirectly* via intrinsically asymmetric intercellular protein complexes and bidirectional feedbacks, or *directly* via different affinities for protein binding in intercellular complexes. Both models behaved similarly in that they required balanced feedback between Fz and Vang to reproduce the expected *fmi*^-^ and double *Vang*^-^ *fz*^-^ clone phenotypes.

Overall, our work suggests that while many binding and feedback regimes can generate polarity and reproduce *fz*^-^ and *Vang*^-^ single clone phenotypes, *Vang*^-^ *fz*^-^ double clones and *fmi*^-^ clones are key simulations required to constrain model parameters to mirror the *in vivo* reality and apparent symmetry in the system.

We suggest that future experimental studies should focus on two questions. First, what are the molecular parameters of intercellular complex formation and do these support formation of intrinsically asymmetric complexes and/or direct signalling between Fz and Vang? Second, what is the molecular nature of the feedback interactions and can at least two opposing feedbacks be identified between Fz-and Vang-containing complexes?

## Experimental Procedures

### Drosophila genetics

Mitotic clones were generated using the *fz*^*P21*^ allele, which is considered to be a null (Jones et al., 1996), using the MARCM system (Lee and Luo, 1999) and *Ubx-FLP* (Emery et al., 2005). Here, *tub-GAL4* and *UAS-GFP* were expressed in every cell, but *tub-GAL80* suppressed expression of *UAS-GFP* in heterozygous or twin-spot tissue only, thus GFP was only observed in *fz*^-^ clone cells. To generate *Vang*^-^ *fz*^-^ double clones, *UAS-Vang-RNAi* transgenes were introduced distal to *fz* on chromosome 3L, such that they also would only be expressed within clones. Strength of Vang knockdown was visualised by staining for Vang. Figures show results from a line from the TRiP collection (HMS01343). However, to control for potential off-target effects, results were confirmed with a non-overlapping independent *pWIZ* line (Bastock and Strutt, 2007). Full genotypes were:

Fig. 6A,B,F,G:

*Ubx-FLP tubGAL4 UAS-nGFP/+; fz*^*P21*^ *FRT80 / tubGAL80 FRT80*

Fig. 6C,D,H,I:

*Ubx-FLP tubGAL4 UAS-nGFP/+; UAS-Vang-RNAi*^*(TRiP)*^ *fz*^*P21*^ *FRT80 / tubGAL80 FRT80*

### Dissection and Immunohistochemistry

White prepupae were collected and aged as appropriate at 25°C. Pupal wings were dissected at either 28 h APF to visualise polarity protein localisation, or at 32.25 h APF for trichomes. Pupal wings were then fixed and stained as previously described (Warrington et al., 2017). Briefly, pupae were fixed for 30-45 minutes at room temperature, prior to dissection of the pupal wing. Wings were transferred into PBS containing 0.2% Triton X-100 (PTX) and 10% normal goat serum to block prior to antibody incubation. Wings were incubated with antibodies overnight at 4°C, and mounted in 10% glycerol, 1xPBS, containing 2.5% DABCO (pH7.5). Primary antibodies for immunostaining were affinity purified rabbit anti-GFP (ab6556, Abcam, UK), affinity-purified rabbit anti-Fz (Bastock and Strutt, 2007), rat anti-Vang (Strutt and Strutt, 2008) and mouse monoclonal anti-Fmi (Flamingo #74, DSHB, (Usui et al., 1999)). Trichomes were stained using Phalloidin conjugated to Alexa-568 (Molecular Probes). Adult wings were dissected from newly eclosed flies and transferred to a 10 μl drop of PTX in a depression slide for imaging.

### Imaging

Fixed pupal wings were imaged on a Nikon A1R GaAsP confocal microscope using a 60x NA1.4 apochromatic lens, with a pixel size of 138 nm, and the pinhole was set to 1.2 AU. 9 Z-slices separated by 150 nm were imaged, and then the 3 brightest slices around junctions were selected and averaged for each channel in ImageJ. Adult wings were imaged on a fluorescence compound microscope to capture trichomes in brightfield and GFP to mark clonal cells. Since brightfield and GFP signals were in different planes, single slices were selected and realigned in Adobe Photoshop.

### Computational modelling

A detailed description of the model equations, assumptions, conditions, and parameters can be found in the Supplemental Text. Briefly, the computational model comprises a set of coupled ordinary differential equations, which encode intercellular binding reactions between core proteins/complexes and intracellular diffusion in a one-dimensional looped line of cells aligned along a proximodistal axis. This set of equations is solved numerically using an explicit Runge-Kutta method. A MATLAB implementation will be deposited in GitHub. Steady state was considered to be achieved once levels of all molecular species remained constant after solving for a sufficiently long time (for example see Fig.S1E,F). Wild-type polarity was defined as higher levels of bound Fz on distal cell edges with higher levels of Vang on proximal cell edges and generally these differences were strong when such bistability occurred.

A brief description of key model parameters is given here:

*Binding rate constants:* Mass action binding rates are proportional to the product of reactants in any reaction. Forward reaction rates are parameterised by rate constants *k*_*1*_ to *k*_*7*_, and reverse reactions by *v*_*1*_ to *v*_*7*_. The relative reaction rates are thus determined by the dissociation constant K_D_ calculated by *v*_*j*_/*k*_*j*_.

*Diffusion:* In a 1D model where each cell has just two compartments, diffusion is modelled by a linear flux between these compartments. The rate of this flux is parameterised by *D* = *μ/L*^2^, where *μ* is the diffusion coefficient, estimated at 0.03 μm^2^s^−1^ (Fischer et al., 2013; Klünder et al., 2013) and *L* is the distance between compartments, estimated at 5 μm.

*Feedback:* Destabilising feedback is modelled by a Hill function of the form *h*(*x*) = 1 + (*V*_max_ −1)[*x*]^*w*^/(*K*^*w*^ + [*x*]^*w*^), where [*x*] is the amount of protein *x* mediating the feedback.

The parameter *V*_max_ determines the strength of the feedback as the maximum fold-change that can be conferred to the off-rate of each reaction. Since feedback can be mediated by either Fz or Vang, strength of feedback is determined by *V*_*max,F*_ or *V*_*max,V*_, respectively. *K* determines the amount of protein *x* required to switch from weak to strong feedback and *w* determines the rate of this switch. We set *K* to 0.5, which is half of the initial concentration of Fz and Vang in each compartment. A higher value of *w* reflects cooperativity such that a small increase in [*x*] can cause large differences in its ability to destabilise complexes. Since we do not have evidence of enzymatic ability in these reactions which may cause such cooperativity, we aimed to maintain *w* as low, and is thus set to 2 in all simulations shown.

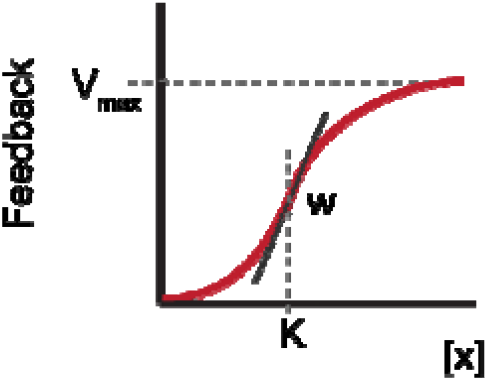

## Acknowledgements

The authors would like to thank Guy Blanchard, Jochen Kursawe, Helen Strutt and Simon Fellgett for comments on the manuscript. We also acknowledge the Bloomington *Drosophila* Stock Center for providing fly strains. The work was funded by a Wellcome Senior Fellowship to DS (WT100986/Z/13/Z), a Vice-Chancellor’s Fellowship from the University of Sheffield to AF and a BBSRC project grant (BB/R016925/1).

## Author Contributions

D.S. conceived and performed experiments, interpreted results, provided expertise and feedback. K.F. designed and wrote the code, ran simulations and interpreted results. A.F. wrote the code, interpreted results and provided expertise and feedback. All authors wrote and edited the manuscript and secured funding.

