## Supplemental_Figures for "Information Flow in Planar Polarity"

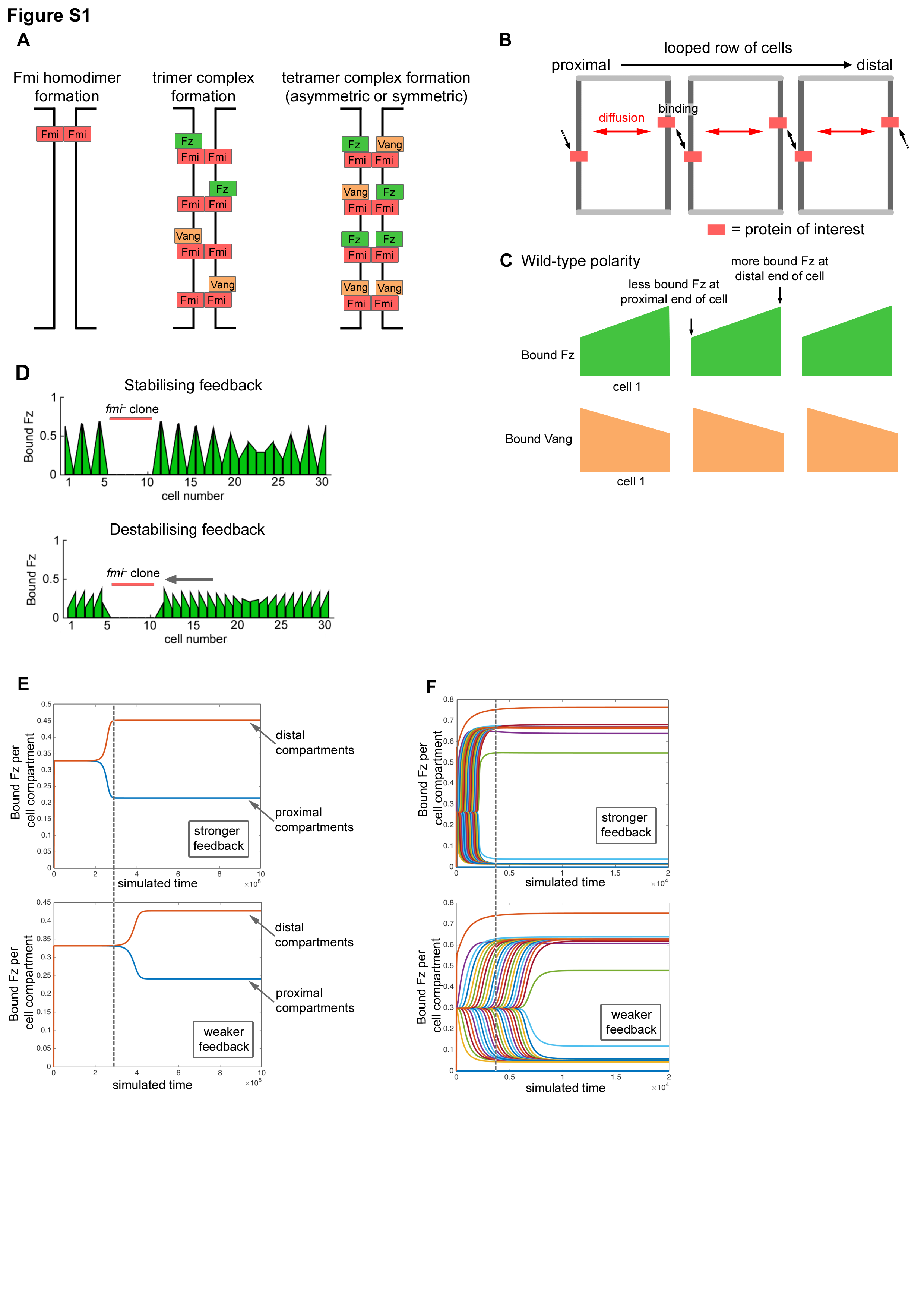

**Figure S1. [Figure 1 – Figure Supplement 1] Model formulation to simulate signalling in planar polarity**.

(A) Complexes form in three stages at the junctions between two neighbouring cells. First, Fmi:Fmi dimers must form, followed by binding of Fz or Vang to either Fmi molecule, generating a trimer. Finally, further binding of either Fz or Vang may occur to form tetrameric asymmetric (Fz and Vang on opposing sides) or symmetric (containing *either* Fz or Vang) complexes.

(B) Polarity is simulated on a one-dimensional row of cells, each with two compartments. We implement periodic boundary conditions such that cells are looped to form a ring. Proteins can localise within these compartments, where they can bind reversibly to form complexes (as in A) or diffuse across the cell.

(C) Wild-type polarity is defined such that Fz, when bound into complexes, accumulates at distal cell ends, whereas Vang accumulates at proximal cell ends. Amounts of bound proteins are plotted to generate a bar for each cell, where a sloped top indicates polarised localisation.

(D) Example simulation result for a *fmi^–^* clone in Model 2 either with only stabilising feedbacks (upper) or only destabilising feedbacks (lower) active from both Fz and Vang (*V*_max,F_  = *V*_max,V_ = 7). When stabilising feedbacks are active, neighbouring cells can adopt opposing polarity (period-two pattern), while when destabilising feedbacks are active, distal non-autonomy is evident with neighbouring cells adopting a common polarity (grey arrow).

(E) Bound Fz in each cell compartment plotted over time from a simulation of Model 2 without any clones, and two destabilising feedback interactions of equal strength. The initial bias drives all distal compartments (overlaid to form orange curve) to have increased levels of bound Fz, compared to proximal compartments (overlaid to form blue curve). Stronger feedback (*V*_max,F_  = *V*_max,V_ = 38) leads to steady state being achieved more quickly than with weaker feedback (*V*_max,F_  = *V*_max,V_ = 32). For comparison, the vertical dashed line indicates the time at which the simulation with stronger feedback reached steady state.

(F) Bound Fz in each cell compartment, plotted over time as differently coloured curves, from simulation of Model 2 with a *fz^–^* clone and no initial bias in Fz localisation. Two destabilising feedback interactions of equal strength are active. Since the polarising signal propagates from the clone boundary, each cell achieves a polarised steady state at a different time, thus curves for individual distal or proximal compartments do not all overlap (as they did for panel E). Stronger feedback (*V*_max,F_  = *V*_max,V_ = 20) leads to steady state being achieved more quickly than with weaker feedback (*V*_max,F_  = *V*_max,V_ = 10). For comparison, the vertical dashed line indicates the time at which the simulation with stronger feedback reached steady state.

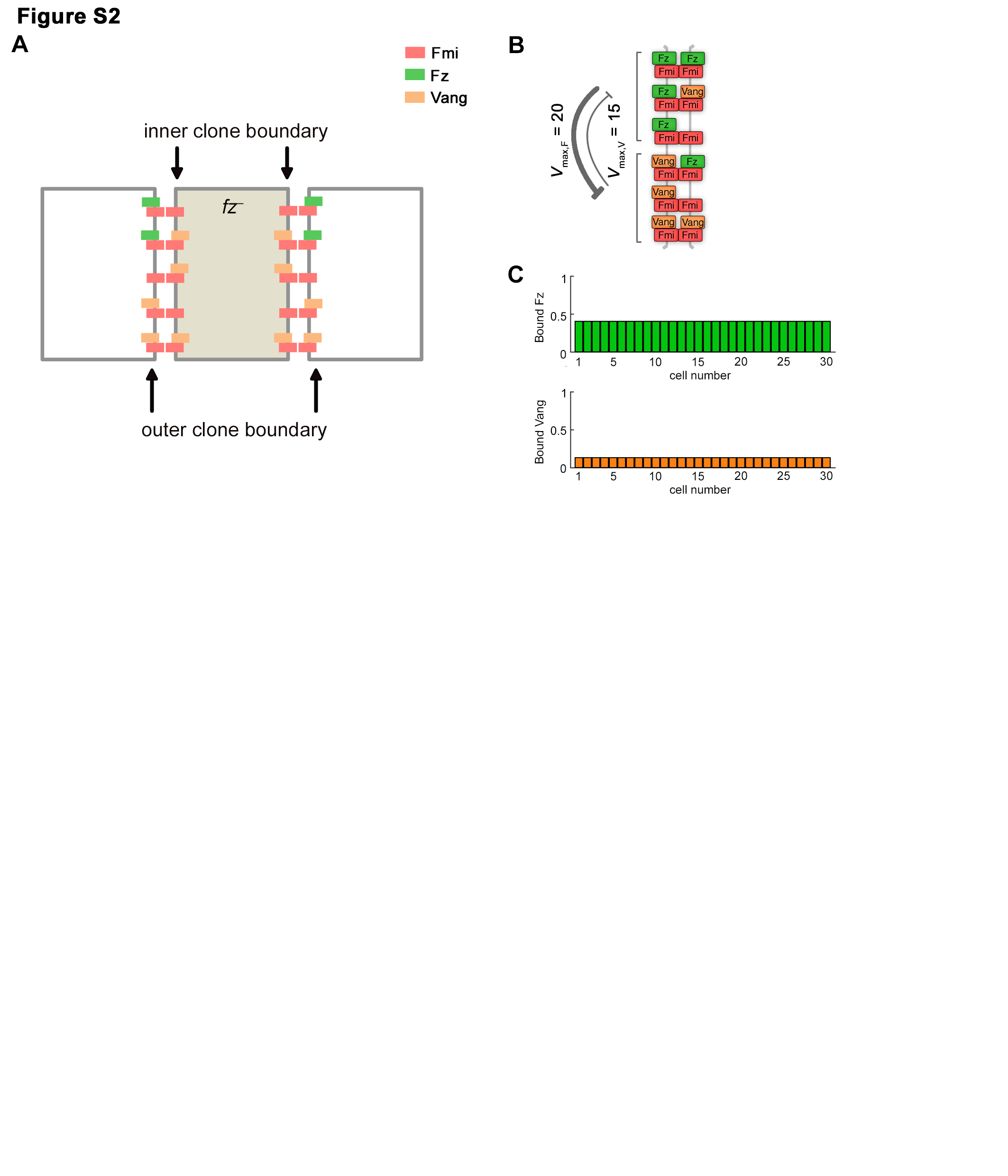

**Figure S2. [Figure 2 – Figure Supplement 1] Model 1 with no direct signalling does not generate non-autonomy around clones.**

(A) Diagram of complex formation with ‘no direct signalling’ at the boundary of a *fz*^–^ clone. There are equal possibilities for both Fz and Vang to bind at the outer clone boundaries. Therefore feedback interactions on this boundary do not favour one molecule over the other and the small initial bias in unbound Fz is the only cue for polarisation of such complexes.

(B) Diagram of feedbacks acting with an example of unbalanced strengths, such that feedback from Fz is stronger than that of Vang.

(C) Simulation of a wild-type field of cells with no direct signalling and unbalanced feedback strengths (*V_max_*_,F_ = 20, *V_max_*_,V_ = 15). This system does not generate a polarised steady state. At steady state there is more bound Fz (upper panel) than bound Vang (lower panel) due to the increased strength of the destabilising feedback from Fz.

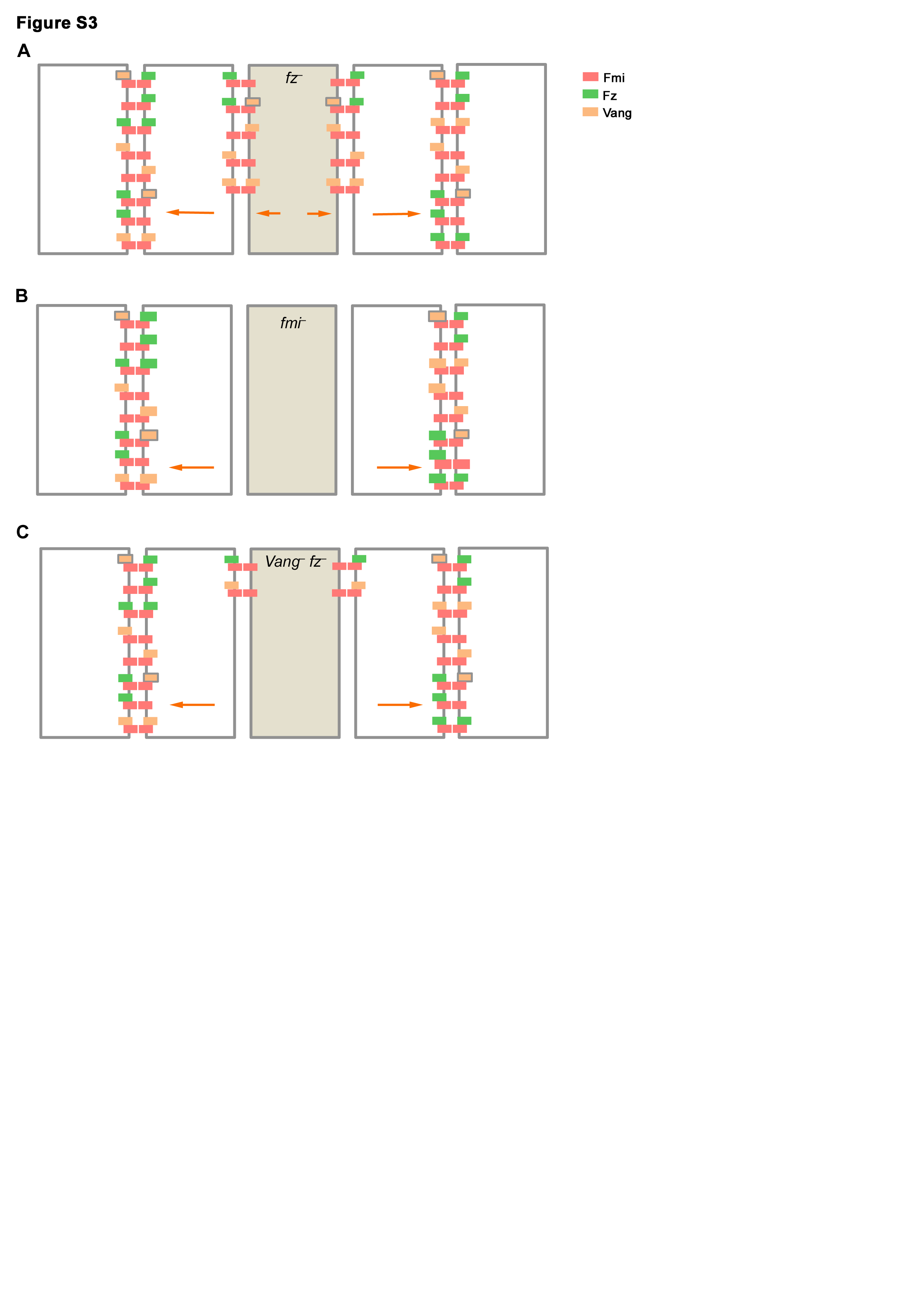

**Figure S3. [Figure 3 – Figure Supplement 1] Complex formation with direct monodirectional signalling**

(A) Diagram of complex formation at the boundary of a *fz*^–^ clone with direct monodirectional signalling. The monodirectional signal results in Vang preferentially binding to complexes that contain Fz (orange boxes with grey outline). Since such complexes cannot form on the outer clone boundary abutting a *fz*^–^ clone, Vang preferentially binds to the edge furthest from the clone. Similarly, Vang in clone cells that neighbour wild-type cells preferentially localises towards wild-type neighbours where it can bind to Fz containing complexes. Orange arrows indicate the preferred direction of Vang localisation in individual cells, caused by the monodirectional signal.

(B) Diagram of complex formation in cells neighbouring a *fmi*^–^ clone. In the neighbouring cells of *fmi*^–^ clones, no complexes can form at the clone boundary, thus all of the Fz and Vang for the cell must localise away from the clone (larger green/orange boxes). Since the monodirectional signal results in Vang preferentially binding to complexes that contain Fz (orange boxes with grey outline), its binding is favoured over on this cell edge. Thus, in cells neighbouring the clone, Vang preferentially localises to cell edges away from the clone (orange arrows), driving polarity direction and generating distal non-autonomy.

(C) Diagram of complex formation in cells neighbouring a *Vang*^–^ *fz*^–^ clone. In the neighbouring cells of *Vang*^–^ *fz*^–^ clones, only trimer complexes can form at the clone boundary. Since the monodirectional signal results in Vang preferentially binding to complexes that contain Fz (orange boxes with grey outline), its binding is favoured in the neighbouring cell on the edge furthest from the clone. Thus, in cells neighbouring the clone, Vang preferentially localises to cell edges away from the clone (orange arrows), driving polarity direction and generating distal non-autonomy.

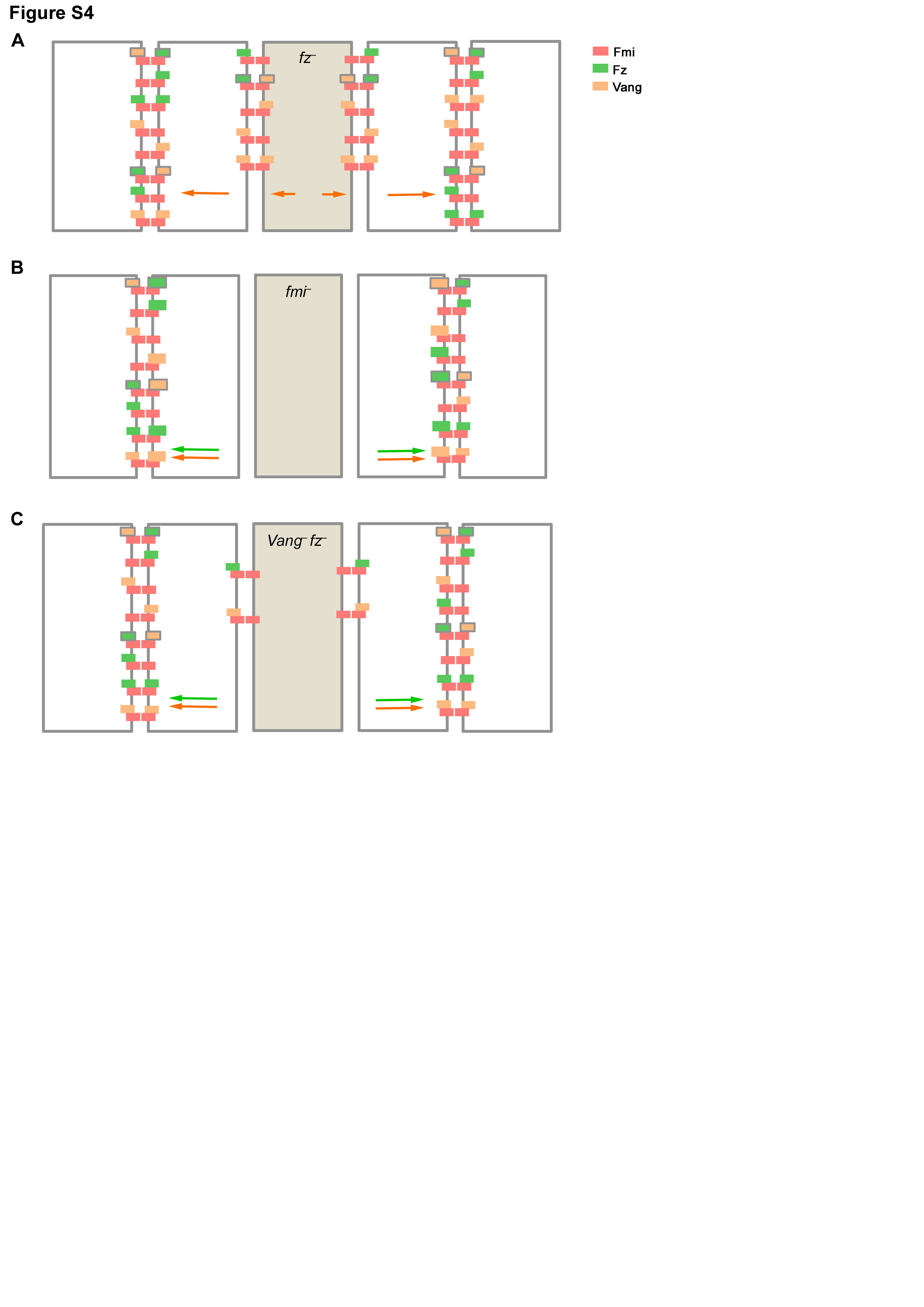

**Figure S4. [Figure 4 – Figure Supplement 1] Complex formation with direct bidirectional signalling**

(A) Diagram of complex formation at the boundary of a *fz*^–^ clone with direct bidirectional signalling. The bidirectional signal results in both Fz and Vang preferentially binding to asymmetric tetramer complexes (i.e. those with green/orange boxes with grey outline). In cells immediately neighbouring a *fz*^–^ clone, these preferred stable complexes can only form in one orientation, thus Vang preferentially binds to the edge furthest from the clone (orange arrows). Similarly, Vang in clone cells that neighbour wild-type cells preferentially localises to cell edges towards the wild-type neighbours.

(B) Diagram of complex formation in cells neighbouring a *fmi*^–^ clone. In the neighbouring cells of *fmi*^–^ clones, no complexes can form at the clone boundary, thus all of the Fz and Vang for the cell must localise away from the clone (indicated by larger green/orange boxes). Both Fz and Vang have lower dissociation constants when in asymmetric tetramer complexes, thus both preferentially localise to cell edges away from the clone (green/orange arrows). Since neither outcompetes the other on this boundary, the next cell polarises normally according to the global cue.

(C) Diagram of complex formation in cells neighbouring a *Vang*^–^ *fz*^–^ clone. In the neighbouring cells of *Vang*^–^ *fz*^–^ clones, only trimeric complexes can form at the clone boundary. Both Fz and Vang have lower dissociation constants when in asymmetric tetramer complexes, thus both preferentially localise to cell edges away from the clone (green/orange arrows). Since neither outcompetes the other on this boundary, the next cell polarises normally according to the global cue.

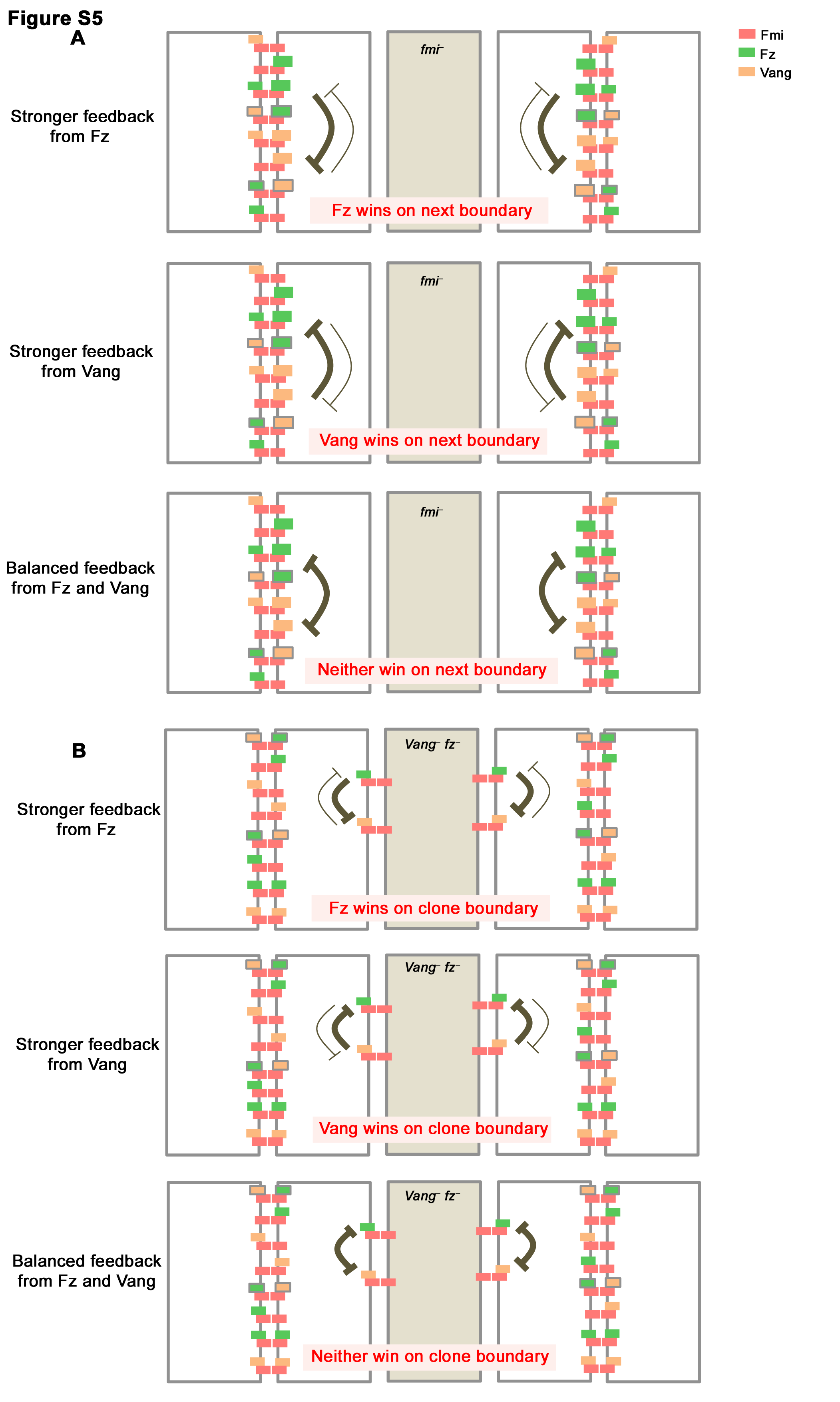

**Figure S5. [Figure 4 – Figure Supplement 2] Unbalanced feedback strengths generate non-autonomy around *fmi*^–^ and *Vang*^–^ *fz*^–^ clones in a direct bidirectional model.**

(A) Diagram of complex formation in cells neighbouring a *fmi*^–^ clone. In the neighbouring cells of *fmi*^–^ clones, no complexes can form at the clone boundary, thus all of the Fz and Vang for the cell must localise away from the clone (larger green/orange boxes). Both Fz and Vang have lower dissociation constants when in asymmetric tetramer complexes (boxes with grey outlines). If there is stronger feedback from Fz (top), Fz outcompetes Vang on these boundaries generating proximal non-autonomy. However, if there is stronger feedback from Vang (middle), Vang outcompetes Fz on these boundaries generating distal non-autonomy. If feedbacks are balanced, clones are autonomous (bottom).

(B) Diagram of complex formation in cells neighbouring a *Vang*^–^ *fz*^–^ clone. In the neighbouring cells of *Vang*^–^ *fz*^–^ clones, only trimeric complexes can form at the clone boundary. If there is stronger feedback from Fz (top), Fz outcompetes Vang on these boundaries generating distal non-autonomy. However, if there is stronger feedback from Vang (middle), Vang outcompetes Fz on these boundaries generating proximal non-autonomy. If feedbacks are balanced, clones are autonomous (bottom).
