## Supplemental_Text for "Information Flow in Planar Polarity"

### Model formulation

#### Tissue geometry and boundary conditions

We model planar polarity complex formation in a one-dimensional row of 30 cells, each having a proximal, or left ($^L$), and distal, or right ($^R$), compartment. For simplicity, we impose periodic boundary conditions such that the cell row is looped to form a ring, and assume that each cell has the same size. Where present, mutant clones are 5 cells wide. Depending on parameter values, these cell numbers are sufficient for us to observe boundary effects in cells near to a clone, but to still observe cells with wild-type polarity away from the clone. Note that as long as a clone is more than one cell wide, its actual size does not alter any non-autonomous effects in our model, since these relate only to the clone boundary.

#### Biochemical reactions

In our model, proteins can localise within cellular compartments, bind to one another in juxtaposed compartments between neighbouring cells, and redistribute to the other compartment within a cell. For simplicity, we consider only the transmembrane proteins Flamingo (Fmi), Frizzled (Fz) and Van Gogh (Vang), since evidence suggests that they are the key components in cell-cell signalling (see main text). We assume that Fmi can form a homodimeric bridge between cells, that Fz and Vang can each bind to Fmi in the same cell compartment, and that once Fz is bound, Vang cannot bind to the same Fmi molecule due to steric hindrance (and vice versa). Thus, the following reversible binding reactions can occur at each cell-cell interface, where $^\dagger$ denotes a protein or complex in a neighbouring cell:

$$\text{Fmi} + \text{Fmi}^\dagger \rightleftharpoons \text{Fmi:Fmi} \tag{1}$$

$$\text{Fz} + \text{Fmi:Fmi}^\dagger \rightleftharpoons \text{Fz-Fmi:Fmi} \tag{2}$$

$$\text{Vang} + \text{Fmi:Fmi}^\dagger \rightleftharpoons \text{Vang-Fmi:Fmi} \tag{3}$$

$$\text{Vang}^\dagger + \text{Fz-Fmi:Fmi} \rightleftharpoons \text{Fz-Fmi:Fmi-Vang} \tag{4}$$

$$\text{Fz} + \text{Vang-Fmi:Fmi}^\dagger \rightleftharpoons \text{Fz-Fmi:Fmi-Vang} \tag{5}$$

$$\text{Vang} + \text{Vang-Fmi:Fmi}^\dagger \rightleftharpoons \text{Vang-Fmi:Fmi-Vang} \tag{6}$$

$$\text{Fz} + \text{Fz-Fmi:Fmi}^\dagger \rightleftharpoons \text{Fz-Fmi:Fmi-Fz} \tag{7}$$

Note that oppositely oriented ($^\dagger$) complexes can also form. The ':' between Fmi molecules indicates binding across the junction of two neighbouring cells.

There is evidence that Fz and Vang can interact directly across cell junctions, possibly stabilising the complex [1, 2]. While we do not explicitly account for this within the complexes that can form, we do explore models where complexes have increased stability when both Fz and Vang are present.

*In vivo* experiments show polarity can arise in approximately 8 hours in the pupal wing [3, 4, 5, 6], during which time protein levels as observed by immunofluorescence do not significantly vary. Therefore, we neglect protein synthesis and degradation, hence the total amounts of Fz, Fmi and Vang are conserved in each cell.

### Governing equations

Based on the above considerations, the amounts of each biochemical species in cell $i$ satisfy the following system of ordinary differential equations:

$$\frac{\mathrm{d}}{\mathrm{d}t}[\mathrm{Fmi}]_i^L = -R_i^{(1)} - R_i^{(10)}, \tag{8}$$
$$\frac{\mathrm{d}}{\mathrm{d}t}[\mathrm{Fmi}]_i^R = -R_{i+1}^{(1)} + R_i^{(10)}, \tag{9}$$
$$\frac{\mathrm{d}}{\mathrm{d}t}[\mathrm{Fz}]_i^L = -R_i^{(2L)} - R_i^{(5L)} - R_i^{(7L)} - R_i^{(8)}, \tag{10}$$
$$\frac{\mathrm{d}}{\mathrm{d}t}[\mathrm{Fz}]_i^R = -R_i^{(2R)} - R_i^{(5R)} - R_{i+1}^{(7R)} + R_i^{(8)}, \tag{11}$$
$$\frac{\mathrm{d}}{\mathrm{d}t}[\mathrm{Vang}]_i^L = -R_i^{(3L)} - R_{i-1}^{(4R)} - R_i^{(6L)} - R_i^{(9)}, \tag{12}$$
$$\frac{\mathrm{d}}{\mathrm{d}t}[\mathrm{Vang}]_i^R = -R_i^{(3R)} - R_{i+1}^{(4L)} - R_{i+1}^{(6R)} + R_i^{(9)}, \tag{13}$$
$$\frac{\mathrm{d}}{\mathrm{d}t}[\mathrm{Fmi{:}Fmi}]_i = R_i^{(1)} - R_i^{(2L)} - R_{i-1}^{(2R)} - R_i^{(3L)} - R_{i-1}^{(3R)}, \tag{14}$$
$$\frac{\mathrm{d}}{\mathrm{d}t}[\mathrm{Fz\text{-}Fmi{:}Fmi}]_i^L = R_i^{(2L)} - R_i^{(4L)} - R_i^{(7R)}, \tag{15}$$
$$\frac{\mathrm{d}}{\mathrm{d}t}[\mathrm{Fz\text{-}Fmi{:}Fmi}]_i^R = R_i^{(2R)} - R_i^{(4R)} - R_{i+1}^{(7L)}, \tag{16}$$
$$\frac{\mathrm{d}}{\mathrm{d}t}[\mathrm{Vang\text{-}Fmi{:}Fmi}]_i^L = R_i^{(3L)} - R_{i-1}^{(5R)} - R_i^{(6R)}, \tag{17}$$
$$\frac{\mathrm{d}}{\mathrm{d}t}[\mathrm{Vang\text{-}Fmi{:}Fmi}]_i^R = R_i^{(3R)} - R_{i+1}^{(5L)} - R_{i+1}^{(6L)}, \tag{18}$$
$$\frac{\mathrm{d}}{\mathrm{d}t}[\mathrm{Fz\text{-}Fmi{:}Fmi\text{-}Vang}]_i^L = R_i^{(4L)} + R_i^{(5L)}, \tag{19}$$
$$\frac{\mathrm{d}}{\mathrm{d}t}[\mathrm{Fz\text{-}Fmi{:}Fmi\text{-}Vang}]_i^R = R_i^{(4R)} + R_i^{(5R)}, \tag{20}$$
$$\frac{\mathrm{d}}{\mathrm{d}t}[\mathrm{Fz\text{-}Fmi{:}Fmi\text{-}Fz}]_i = R_i^{(7L)} + R_i^{(7R)}, \tag{21}$$
$$\frac{\mathrm{d}}{\mathrm{d}t}[\mathrm{Vang\text{-}Fmi{:}Fmi\text{-}Vang}]_i = R_i^{(6L)} + R_i^{(6R)}, \tag{22}$$

where, assuming mass action kinetics (with parameters for binding rate constants ($k_1, \ldots, k_7$) and unbinding rate constants ($v_1, \ldots, v_7$) and simple diffusion (parameterised by $D$), the reaction rates $R_i^{(1)}, \ldots, R_i^{(10)}$ are given by:

$$R_i^{(1)} = k_1[\text{Fmi}]_i^L[\text{Fmi}]_{i-1}^R - v_1[\text{Fmi:Fmi}]_i, \tag{23}$$

$$R_i^{(2L)} = k_2[\text{Fz}]_i^L[\text{Fmi:Fmi}]_i - v_2 h_V\Big([\text{Bd-Vang}]_i^L\Big)[\text{Fz-Fmi:Fmi}]_i^L, \tag{24}$$

$$R_i^{(2R)} = k_2[\text{Fz}]_i^R[\text{Fmi:Fmi}]_{i+1} - v_2 h_V\Big([\text{Bd-Vang}]_i^R\Big)[\text{Fz-Fmi:Fmi}]_i^R, \tag{25}$$

$$R_i^{(3L)} = k_3[\text{Vang}]_i^L[\text{Fmi:Fmi}]_i - v_3 h_F\Big([\text{Bd-Fz}]_i^L\Big)[\text{Vang-Fmi:Fmi}]_i^L, \tag{26}$$

$$R_i^{(3R)} = k_3[\text{Vang}]_i^R[\text{Fmi:Fmi}]_{i+1} - v_3 h_F\Big([\text{Bd-Fz}]_i^R\Big)[\text{Vang-Fmi:Fmi}]_i^R, \tag{27}$$

$$R_i^{(4L)} = k_4[\text{Vang}]_{i-1}^R[\text{Fz-Fmi:Fmi}]_i^L - v_4 h_F\Big([\text{Bd-Fz}]_{i-1}^R\Big)[\text{Fz-Fmi:Fmi-Vang}]_i^L, \tag{28}$$

$$R_i^{(4R)} = k_4[\text{Vang}]_{i+1}^L[\text{Fz-Fmi:Fmi}]_i^R - v_4 h_F\Big([\text{Bd-Fz}]_{i+1}^L\Big)[\text{Fz-Fmi:Fmi-Vang}]_i^R, \tag{29}$$

$$R_i^{(5L)} = k_5[\text{Fz}]_i^L[\text{Vang-Fmi:Fmi}]_{i-1}^R - v_5 h_V\Big([\text{Bd-Vang}]_i^L\Big)[\text{Fz-Fmi:Fmi-Vang}]_i^L, \tag{30}$$

$$R_i^{(5R)} = k_5[\text{Fz}]_i^R[\text{Vang-Fmi:Fmi}]_{i+1}^L - v_5 h_V\Big([\text{Bd-Vang}]_i^R\Big)[\text{Fz-Fmi:Fmi-Vang}]_i^R, \tag{31}$$

$$R_i^{(6L)} = k_6[\text{Vang}]_i^L[\text{Vang-Fmi:Fmi}]_{i-1}^R - v_6 h_F\Big([\text{Bd-Fz}]_i^L\Big)[\text{Vang-Fmi:Fmi-Vang}]_i, \tag{32}$$

$$R_i^{(6R)} = k_6[\text{Vang}]_{i-1}^R[\text{Vang-Fmi:Fmi}]_i^L - v_6 h_F\Big([\text{Bd-Fz}]_{i-1}^R\Big)[\text{Vang-Fmi:Fmi-Vang}]_i, \tag{33}$$

$$R_i^{(7L)} = k_7[\text{Fz}]_i^L[\text{Fz-Fmi:Fmi}]_{i-1}^R - v_7 h_V\Big([\text{Bd-Vang}]_i^L\Big)[\text{Fz-Fmi:Fmi-Fz}]_i, \tag{34}$$

$$R_i^{(7R)} = k_7[\text{Fz}]_{i-1}^R[\text{Fz-Fmi:Fmi}]_i^L - v_7 h_V\Big([\text{Bd-Vang}]_{i-1}^R\Big)[\text{Fz-Fmi:Fmi-Fz}]_i, \tag{35}$$

$$R_i^{(8)} = D\Big([\text{Fz}]_i^L - [\text{Fz}]_i^R\Big), \tag{36}$$

$$R_i^{(9)} = D\Big([\text{Vang}]_i^L - [\text{Vang}]_i^R\Big), \tag{37}$$

$$R_i^{(10)} = D\Big([\text{Fmi}]_i^L - [\text{Fmi}]_i^R\Big), \tag{38}$$

where we have introduced the shorthand notation:

$$[\text{Bd-Fz}]_i^L = [\text{Fz-Fmi:Fmi}]_i^L + [\text{Fz-Fmi:Fmi-Fz}]_i + [\text{Fz-Fmi:Fmi-Vang}]_i^L, \tag{39}$$

$$[\text{Bd-Fz}]_i^R = [\text{Fz-Fmi:Fmi}]_i^R + [\text{Fz-Fmi:Fmi-Fz}]_{i+1} + [\text{Fz-Fmi:Fmi-Vang}]_i^R, \tag{40}$$

$$[\text{Bd-Vang}]_i^L = [\text{Vang-Fmi:Fmi}]_i^L + [\text{Vang-Fmi:Fmi-Vang}]_i + [\text{Fz-Fmi:Fmi-Vang}]_{i-1}^R, \tag{41}$$

$$[\text{Bd-Vang}]_i^R = [\text{Vang-Fmi:Fmi}]_i^R + [\text{Vang-Fmi:Fmi-Vang}]_{i+1} + [\text{Fz-Fmi:Fmi-Vang}]_{i+1}^L. \tag{42}$$

To generate a bistable system where polarity can be stable in either proximal or distal direction, we introduce regulation in the form of locally destabilising feedback interactions, represented in equations (24)–(35) by Hill functions of the form:

$$h(x) = 1 + \frac{(V_{\max,x} - 1)[x]^w}{K^w + [x]^w}, \tag{43}$$

Here $[x]$ denotes the concentration of protein $x$ (Bd-Fz or Bd-Vang) causing the feedback. The parameter $V_{\max,x}$ determines the strength of the feedback as the maximum fold-change that can be conferred to the off-rate of each reaction. The parameter $K$ determines the concentration of $x$ required to switch from weak to strong feedback and $w$ determines the rate of this switch.

We also test non-autonomous phenotypes when using stabilising feedback interactions. These are of the same form as in equation (43), but are used to regulate on-rates rather than off-rates.

Reactions $R_i^{(8)}$, $R_i^{(9)}$, $R_i^{(10)}$ represent diffusion of each unbound molecule within cells. For simplicity, we assume that Fz, Vang and Fmi share a common diffusion constant, $D$. Since our aim is to explore the qualitative, rather than quantitative, behaviours of this model, all biochemical species are assumed to have arbitrary units.

### Initial conditions

Each cell is initialised with two arbitrary units of Fz, Vang and Fmi. For Vang and Fmi, these are equally distributed among compartments. Although the upstream cue to generate cellular asymmetry of complexes is unknown, it has been demonstrated that Fz may be trafficked in a distal direction along microtubules [7]; we therefore assume that a small proportion of cellular Fz (an initial bias; $b = 0.001$) is localised to the distal compartment of each cell, resulting in 0.999 units proximally localised and 1.001 units localised distally in each cell. This provides an initial polarity cue, which can then amplified by feedback interactions.

It should be noted that altering the magnitude of this initial bias ($b$) did not alter the direction of non-autonomy observed around clones, but did affect the range of the non-autonomy.

### Parameter values

The parameters in our model are binding rate constants ($k_1, \ldots, k_7$), unbinding rate constants ($v_1, \ldots, v_7$), feedback parameters ($V_{\max,F}$, $V_{\max,V}$, $K$, $w$), and a diffusion constant ($D$). We set binding rate constants $k_1, \ldots, k_5$ to 1 in all simulations. In Model 4 where only asymmetric complexes are modelled binding rate constants $k_6$ and $k_7$ are zero such that symmetric complexes cannot form. In all other where symmetric complexes were allowed to form (Models 1, 2 and 3), $k_6$ and $k_7$ were equal to 1.

We consider different values of unbinding rate constants depending on the specific model, as depicted via the dissociation constant ($K_D = v_j/k_j$) in Figures 2, 3, 4 and 5. These values are chosen such that Fmi:Fmi dimers have a much higher off-rate than other complexes; an assumption consistent with experimental evidence showing that Fmi is lost from cell-cell junctions in the absence of Fz and Vang [1, 4].

We take $D = \mu/L^2$, where the diffusion coefficient $\mu = 0.03\ \mu\text{m}^2\text{s}^{-1}$ and $L = 5\ \mu$m is the width of each cell [8, 9].

For wild-type simulations we find that varying $D$ over two orders of magnitude does not alter the stable steady state, only the timescale over which it is reached (not shown). For simulations with clones, we find that faster diffusion can result in small changes in the degree of propagation, but not its direction.

In equation (43), $K$ determines the amount of $x$ required to switch from weak to strong feedback. In all simulations we set $K$ to 0.5, which is half of the initial concentration of Fz and Vang in each compartment. For each destabilising feedback interaction, the value of the maximum fold-difference in off-rate, $V_{\mathrm{max}}$, is presented in the relevant Figure, but in general is maintained below 10. For simulations where only one feedback is included, the other is switched off by setting its $V_{\mathrm{max}}$ value to 1. For all simulations, we set the Hill coefficient $w$ to 2, reflecting our assumption that cooperativity is low since these interactions are primarily thought to arise from steric hindrance rather than enzymatic activity.

#### Numerical solution

The set of coupled ordinary differential equations is solved numerically using an explicit Runge-Kutta method. Simulations were run to ensure that steady state was achieved by plotting solutions over time to ensure no further change in levels of molecular species (see Fig.S1E,F). A MATLAB implementation will be deposited in GitHub.

### Feedback interactions: stabilising or destabilising?

As described in the main text, both stabilising and destabilising interactions have been suggested as mechanisms for generating a bistable polarity system (Fig.1G), thus we analysed both in our models. We found that each type of feedback produced similar phenotypes around clones; however, in certain cases there were subtle differences. The greatest difference was that in cases where only stabilising feedback was present, there was no mechanism preventing complexes of opposite orientations accumulating on the same cell-cell junction. In some cases this led to propagating period-two patterns where every other cell adopts the same polarity and thus, neighbouring cells have opposing polarity (Fig.S1D, upper panel), which does not occur when using destabilising feedbacks alone (Fig.S1D, lower panel). We found that for stabilsing feedback between like complexes, the system is slower to polarise due to decreased sorting of complexes to appropriate ends of the cell, and thus more sensitive to the rate of diffusion. In the simulations presented within this manuscript we examined models with destabilising feedbacks only.

### Range of non-autonomy

As model parameters vary, so does the range of non-autonomy observed around clones. In this section, we address what controls this range. In each model every cell is capable of responding to a polarity cue. In a wild-type simulation, the only cue that each cell receives is the small initial distal bias in Fz localisation ($b$). The feedback interactions then act to amplify that initial bias until the stable polarised steady state is achieved. The rate at which steady state is reached depends primarily on the feedback strength, controlled by parameters $V_{\max,F}$ and $V_{\max,V}$, although if the initial bias is larger, steady state will be achieved more rapidly.

For example, in the case of destabilising feedback, if feedback is weak, complexes have only weak effects on one another. Thus, complexes are slow to sort to a polarised steady state. However, if feedback is strong then complexes of opposite orientations have a strong ability to destabilise one another. Sorting of complexes is much faster and thus the polarised steady state is achieved rapidly (Fig.S1E).

In a clone scenario, but in the absence of the initial bias in Fz localisation, there is an alternative polarity cue in the system in the form of the clone boundary. In the case of a $fz^-$ clone, cells immediately neighbouring the clone polarise according to this boundary signal (see main text for rationalisation of complex formation at clone boundaries). This boundary signal is then propagated from cell to cell in both proximal and distal directions. The speed of propagation depends on the strength of the feedback, with stronger feedback leading to faster propagation (Fig.S1F).

In simulations with both a clone *and* an initial bias, cells must polarise according to the two cues, which may be in opposing directions. For example, in cells neighbouring the distal side of a $fz^-$ clone, there is a boundary cue recruiting Fz complexes to the proximal cell edge, competing with the initial distal bias ($b$) in unbound Fz within the cell. When feedback is weak, the initial distal bias in Fz localisation is amplified slowly and although propagation from the clone is also slower, more cells show non-autonomy. Alternatively, when feedback is strong, wild-type cells rapidly polarise and are resistant to reversals propagating from the clone.
